# Neurons underlying aggressive actions that are shared by both males and females in *Drosophila*

**DOI:** 10.1101/2024.02.26.582148

**Authors:** Liangyu Tao, Deven Ayembem, Victor J. Barranca, Vikas Bhandawat

**Author notes:** Correspondence to Vikas Bhandawat.

## Abstract

Aggression involves both sexually monomorphic and dimorphic actions. How the brain implements these two types of actions is poorly understood. We found that a set of neurons, which we call CL062, previously shown to mediate male aggression also mediate female aggression. These neurons elicit aggression acutely and without the presence of a target. Although the same set of actions is elicited in males and females, the overall behavior is sexually dimorphic. The CL062 neurons do not express *fruitless*, a gene required for sexual dimorphism in flies, and expressed by most other neurons important for controlling fly aggression. Connectomic analysis suggests that these neurons have limited connections with *fruitless* expressing neurons that have been shown to be important for aggression, and signal to different descending neurons. Thus, CL062 is part of a monomorphic circuit for aggression that functions parallel to the known dimorphic circuits.

## Introduction

Most animals display sexually dimorphic behaviors (Darwin, 1874). One such behavior is aggression, which is important for defending and obtaining different resources necessary for survival and reproduction (Lorenz, 1963). Circuits underlying aggression have been studied in many organisms, including primates, rodents, and flies (Hoopfer, 2016; Lischinsky and Lin, 2020; Nelson and Trainor, 2007). Perhaps due to the high degree of behavioral dimorphism, much of what we know about the neural circuitry underlying aggression has been studied in a sex-specific manner (Asahina, 2018; Dulac and Kimchi, 2007; Manoli et al., 2013; Yamamoto and Koganezawa, 2013; Yang and Shah, 2014). Yet, aggression is not fully dimorphic, as both vertebrates (Scott, 1966) and invertebrates (Nilsen et al., 2004) show monomorphic (shared by both sexes) motor actions during aggression (Hashikawa et al., 2018; Pandolfi et al., 2021). How neural circuits are organized to drive these shared aspects of aggression is largely unknown.

Like other animals, *Drosophila* exhibits both sexually dimorphic and shared actions during aggression. Sexually dimorphic actions include male-specific lunging and boxing as well as female headbutting and shoving (Chen et al., 2002; Nilsen et al., 2004; Sturtevant, 1915); sexually shared actions include wing threats, charging, approach, standing still, and fencing actions during aggression. While these actions are shared, there is still sexual dimorphism in the details of how they are deployed (Vrontou et al., 2006). For instance, female wing threats are of shorter duration than those of males (Nilsen et al., 2004). Due to the dimorphism in action implementation despite the commonality in the action itself, it is unknown if these actions are driven by sexually dimorphic neurons and circuits or through a common set of neurons within a potentially dimorphic circuit.

Much of the work dissecting the neural circuits underlying fly aggression has focused on dimorphic neurons with a particular focus on neurons involved in male aggression. Most of these neurons express the sex-determination genes *fruitless*(*fru*) and/or *doublesex*(*dsx*) (Koganezawa et al., 2016; Siwicki and Kravitz, 2009; Vrontou et al., 2006; Wohl et al., 2020). In particular, the P1a cluster of *fru*^+^ and *dsx^+^*neurons plays a role in promoting both male mating and male aggression (Anderson, 2016). Neurons in this cluster function by inducing a persistent intrinsic state, lasting for minutes (Hoopfer et al., 2015), which in the presence of another male fly leads to persistent aggression. The neurons that carry signals downstream of P1a are themselves *fru*^+^ (Jung et al., 2020b). Other neurons, all dimorphic, have been found to enhance actions due to aggression through a variety of neurotransmitters, including tachykinin (Asahina et al., 2014), drosulfakinin (Wu et al., 2020), and octopamine (Watanabe et al., 2017). Recently another set of neurons that are not *fru*^+^ have been found to elicit aggressive behaviors (Duistermars et al., 2018); these neurons appear to be functionally unrelated to the *fru*^+^ population (Duistermars et al., 2018). The *fru*^+^/*dsx^+^*population that contains the P1a neurons is also important for aggression in females. Females lack P1a neurons, but have other neurons within this cluster called pC1; one subset in this cluster called the pC1d (which are *dsx*^+^) elicits aggressive behaviors in females through its strong and recurrent connections with aIPg neurons (which are *fru*^+^) and together drive female-specific aggressive behavior including head-butting and shoving (Chiu et al., 2021; Deutsch et al., 2020; Hui et al., 2023; Palavicino-Maggio et al., 2019; Schretter et al., 2020). Thus, much progress has been made in understanding the basis of aggression that is dimorphic. In contrast, relatively little is known about monomorphic aggression circuits in *Drosophila*. A recent study identified a set of sexually conserved neurons (Chiu et al., 2021), but these neurons mediate aggressive approaches rather than the action themselves and the aggressive actions elicited downstream occur through the same sexually dimorphic circuits described above.

Here, we report that a set of monomorphic neurons that, upon optogenetic activation, elicit a common set of aggressive actions in both male and female flies. In this study, we aim to answer three questions – what actions, which neurons, and how do these neurons connect to other neurons involved in aggression? To answer the first question, we quantified the temporal progression of actions. We show that despite driving a common set of actions, there is sexual dimorphism in the temporal progression of each action. Second, we developed an experimental preparation that allows us to drive aggressive actions through spatially targeted optogenetics. Using this setup, we identify the neurons responsible for driving these behaviors. Third, we utilize an EM dataset to show that different subsets of these neurons connect to different descending neurons (DNs) in a modular manner, which suggests the presence of parallel descending motor pathways in driving actions. These neurons also appear to have only sparse connections with the previously reported dimorphic neurons and therefore likely function independently.

## Results

### Activation of L320 neurons drives multiple aggressive behaviors in isolated male and female flies

In a behavioral screen on the collection of Janelia split-Gal4 collection of genetic lines (Dolan et al., 2019), we found that optogenetic activation of neurons using the red-shifted channelrhodopsin, CsChrimson, (Klapoetke et al., 2014) in a split-Gal4 line, L320 (GMR33E02-AD ⋂ GMR47B03-DBD) (**Figure 1A**), drives multiple aggressive actions including wing threat, wing extension, thrusting, as well as holding an alert posture in both isolated male and female flies (**Figure 1B and SVideos 1 and 2**). A hallmark of aggressive behavior – seen here (**Figure 1B**, leftmost panel) - is that the wing movement is upwards (**Figure 1B**) as opposed to the sideways wing extension observed during courtship. Aggressive phenotypes elicited by optogenetic activation of the L320-split line can occur in isolation (i.e., without either the presence of any target of their aggression or resources that they typically fight over); this expression of aggression in the absence of a target is rare as most studies report aggressive actions only in the presence of a conspecific (Certel and Kravitz, 2012; Dankert et al., 2009; Kravitz and Fernandez, 2015). To quantify the behavior, we tracked five body parts of the fly - head, thorax, abdomen, and left/right wingtips – on each of the two camera views. The tracked body parts were triangulated to yield five features (**Figure 1C**). The relationship between the observables obtained from these features and the four aggressive actions – wing threat, wing extension, thrusting, and alert stance – is shown in **Figure 1C** and described below.

**Figure 1.**
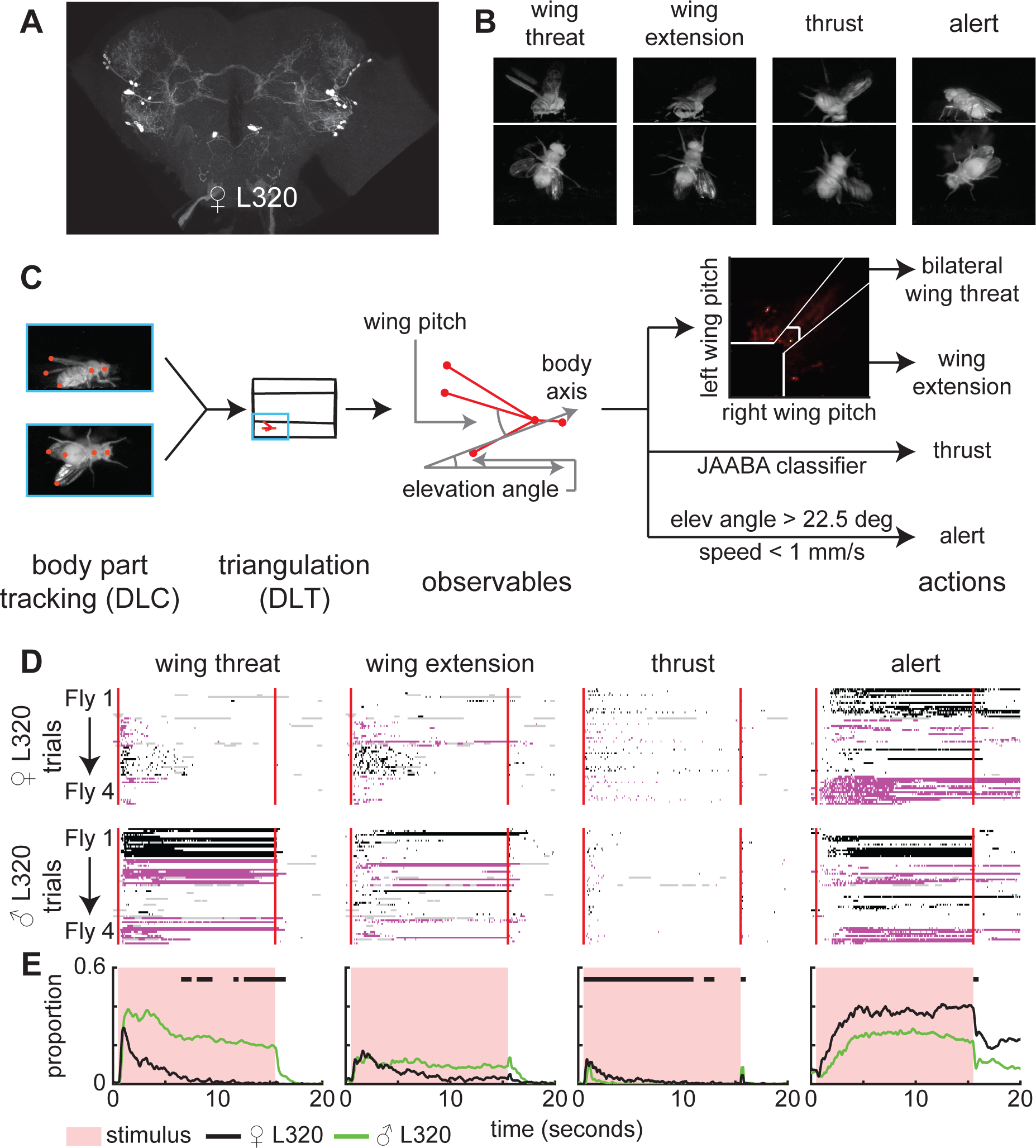
Activation of a small number of neurons drives sexually conserved set of aggressive actions and observables in the absence of sensory information. There is sexual dimorphism in the level of these actions. **A.** Maximum intensity projection (MIP) image of the female L320 brain (L320 > UASChrimson). **B.** Example images of female flies displaying wing threat, wing extension, thrusting, and holding an alert posture. **C.** Analysis pipeline. The head, thorax, abdomen, and two wingtips were tracked for two views using DeepLabCut (DLC). These body parts were 3D triangulated using direct linear transform (DLT). 4 observables (left/right wing pitch, elevation angle, speed) were calculated based on tracked body parts. Wing threat, wing extension, and alert stance were assigned based on thresholded motor elements. Thrusting was found using JAABA. (See methods and Figures 1S1 and 1S3). **D. Top rows:** Single trial ethogram of actions for 4 sample male and female L320 > UASChrimson flies. Trials grouped by flies (fly 1 = trials 1-15, fly 2 = trials 16-30, etc). Red line indicates start and end of optogenetic stimulation period, which lasts for 15 seconds. Alternating black and purple blocks of trials represent different flies. Gray areas indicate >500 ms bouts where there is low confidence in any of the tracked body parts that is used to identify the action. (see methods). **E:** Proportion of all male and female trials where flies are performing each action. Black bars show time points where there is a significant difference between male and female flies (Wilcoxon rank sum test p<0.01, see methods). An 8.9 mW/cm² 617 nm light is turned on during the stimulus period. (Retinal fed: n=12 female flies, n=11 male flies; 15 trials/-fly).

Based on previous work (Nilsen et al., 2004), we defined bilateral wing threat when both wings were raised above 45°. We also found another aggressive action that was distinct from wing threat which we call wing extension in this study: During wing extension, the flies raised one or both wings, but the wing pitch was less than 45° and higher than that found during the control period. Unlike wing threats – which last about one second – wing extension can last much longer and, therefore, we considered it as a separate behavior; the method used to distinguish between wing threat and wing extension is further described in **Figure 1-S1A-C**. We also show that the change in wing pitch is described by two separate behaviors – wing threat and wing extension – with the contribution of wing extension increasing with time. During bilateral wing threats and most wing extensions, flies raised their wings at a ∼45-degree elevation from the body plane; this suggests that these actions represent different forms of wing threat rather than horizontal wing extension exhibited during courtship (**Figure 1-S1E**). While we also observed that many flies will keep their wings slightly ajar when not performing wing threat or extension (**SVideos 1 and 2 and Figure 1-S1C/D**), we focused on wing threat and wing extension in this study.

To identify thrusting, we trained a JAABA classifier using movement speed and elevation angle. Both speed and elevation angle are employed because, at the start of a thrust, flies will lift their front leg and pitch up resulting in an increase in elevation angle and speed. Afterward, flies will push forward and snap down on their front legs often resulting in a lower-than-baseline elevation angle (**Figure 1 S2**). As in previous work (Nilsen et al., 2004), different thrust episodes varied in how the body is elevated and whether only the front legs are lifted off the ground or the other legs as well. There are also differences in how the body snaps back. Our classifier groups different forms of thrusting in a manner comparable to similar grouping together of thrust in previous studies (Nilsen et al., 2004). It is important to note that a similar action – where the fly raises the front part of its body and lunges down – in males is referred to in some studies as a lunge (Hoffmann, 1987). There is some ambiguity in how different authors define these actions (Asahina, 2017); we are following the convention outlined in Nilsen and Kravitz (2004) (Nilsen et al., 2004). Finally, we used a combination of speed and elevation angle thresholds to capture an alert stance when the flies stood still with their front legs straightened in an upward-pitched position (**Figure 1C**). An alert stance is a novel form of aggression that has not been reported, to our knowledge. We observe an alert pose because most work on aggression uses a single camera view which is insufficient for detecting an alert pose. We also observed other behaviors such as retreat and take-off but have not quantified these behaviors in this study. Aggression is characterized by a complex structure consisting of recurring behavioral sequences (Chen et al., 2002; Hoopfer, 2016; Nilsen et al., 2004); we observe a similar complex behavioral structure when activating the L320 neurons (**Figure 1D and Figure 1 S3**). Aggressive actions are not observed in flies that are not fed retinal (**Figure 1-S4**). Ethograms of the first 4.5 seconds are shown in **Figure 1-S6A** and illustrate that there is an alternation between wing threat, wing extension, thrust, and alert state (**Figure 1-S6A**). In most trials, wing threat is the first action and can occur in under 300 millisecond (**Figure 1-S6B**), followed by thrust. Wing extensions and alert posture occurred with more delay. There is considerable variability in the occurrence, latency, and persistence of each of the actions (four males and females are shown in **Figure 1D**, all flies in **Figures 1-S4 and 1-S6**). As an example, wing threat does not occur in any trial in one fly in our dataset (**Figure 1D**) and occurs in only some trials in another fly (**Figure 1D**).

Despite the moment-by-moment action being different across different trials and flies, the probability of observing a given action has a clear temporal progression. The probability of observing wing threat was highest shortly after stimulus onset, decreased rapidly in females and slowly decreasing in males (**Figure 1E**); many males did not show any habituation (**Figures 1E and 1S3**). Wing extension follows a similar trend to wing threat with males showing more persistence (**Figure 1E**). Unlike wing threat and extension, males perform significantly fewer thrusts and exhibit faster habituation of this action (**Figure 1E**). Wing-driven behavior appears to occur a greater proportion of the time compared to thrusts. However, this lower moment-by-moment proportion of flies performing thrusts is due to their transient nature (**Figure 1E**). At light offset, flies will sometimes perform thrusts or controlled jumps that are time-locked to stimulus-off (**Figure 1E**). This results in an instantaneous peak in the proportion of flies performing thrusts (**Figure 1E**). After a period of thrusting, flies will often transition into an alert posture. Unlike other actions, this alert stance is persistent even after the stimulus has been turned off.

#### L320 labels four populations of neurons and a descending neuron

L320-split, despite being sparse, labels multiple populations of neurons (**Figure 1A**). To identify these neurons, we utilized a morphology-matching algorithm called NBLAST to compare traced neurites of L320-split neurons with neuron skeletons within a recently released female electron microscopy dataset called the hemibrain dataset (**Figure 2-S1A and methods**) (Costa et al., 2016; Scheffer et al., 2020). Because the neurite of different neuronal populations intersect, we used multicolor flip (MCFO) to stochastically label 1-3 L320 neurons in each fly brain so that we could trace them unambiguously (**Figure 2-S1B** for one example) (Nern et al., 2015). The traced neurons were morphologically located in the lateral horn (LH), anterior ventrolateral protocerebrum (AVLP), posterior ventrolateral protocerebrum regions (PVLP), as well as a single pair of descending neurons (DN). We found that the anterior cluster of LH neurons, which formed a characteristic u-shaped neurite tract that first runs in the anterior-ventral direction before looping back in the posterior-dorsal direction and arborizes within the lateral portion of the lateral horn, has a similar morphology to LHAV4a in the hemibrain (**Figure 2 and Figure 2-S1C**). LHAV4a are structurally connected to projection neurons implicated in bilateral contrast sensing of cis-Vaccenyl Acetate (cVA) (Taisz et al., 2023). cVA is the primary pheromone implicated in fly courtship and aggression (Kurtovic et al., 2007; Liu et al., 2011; Wang and Anderson, 2010). The posterior cluster of LH neurons was identified as LHPV6a1/3 in the hemibrain. The AVLP neurons have a similar morphology to the CL062 neurons since they both cross the midline along an anterior-posterior curve and form characteristic dorsal-ventral branches. These CL062 neurons have previously been shown to drive aggressive male threats (Duistermars et al., 2018); they are also referred to as AVLP_pr12 neurons(Baker et al., 2022). The PVLP neurons have a similar morphology to PVLP077/078 neurons due to their anterior-ventral to posterior-dorsal neurite track. Finally, none of the descending neurons in the hemibrain dataset matched the DN labeled by L320-split. We did not further search for the DN in the flywire dataset.

**Figure 2.**
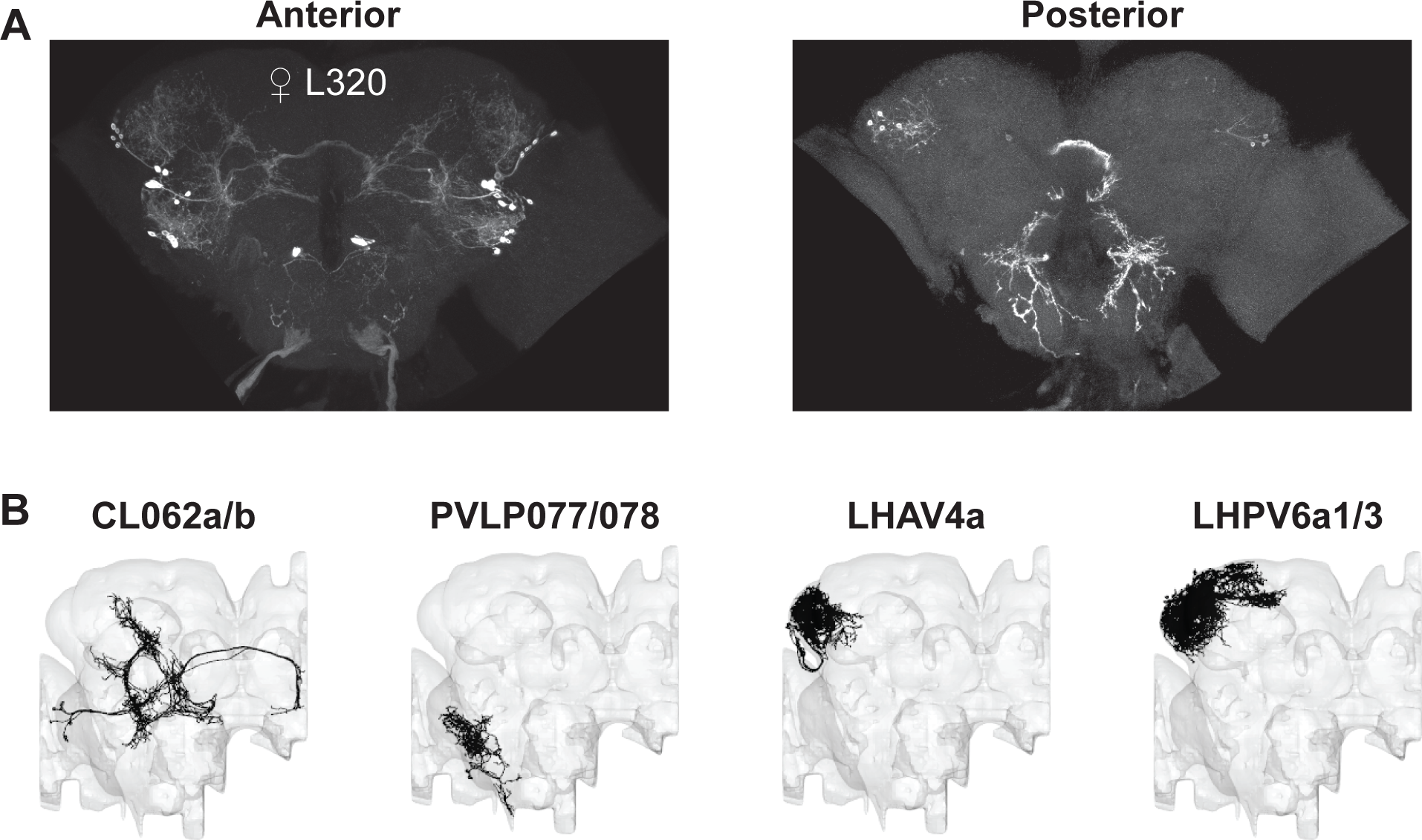
Neuron types labeled by L320. **A.** Maximum projection image of female L320 in the anterior plane (left) and posterior plane (right). L320 labels two types of lateral horn (LH) neurons, one type of anterior ventrolateral protocerebrum neuron (AVLP), one type of posterior VLP (PVLP) neuron, and a pair of descending neurons (DN). **B.** Neuron types found in the hemibrain EM dataset corresponding to each L320 neurons. The DN was not found in the hemi-brain.

#### Activation of CL062 neurons drives aggressive actions

To determine which population of neurons is important for driving aggression, we used a Digital Micromirror Device (DMD) projector to optogenetically activate each cluster of neurons independently (**Figure 3A and Figure 3-S3A**). Since in this setup, flies are head-fixed and walk on an air-supported ball, and aggressive behaviors have never been demonstrated in a head-fixed preparation, we first characterized the effect of activating all L320 neurons. Using light delivered at the image plane, both male and female flies elicited a robust wing response; as in the case of freely walking flies, the wing response was characterized by a change in wing pitch (**Figure 3B**). Perhaps due to variability from one preparation to another, the baseline wing pitch was different across flies (**Figure 3-S1A, SVideo 3**) and led to different physical space available for each wing which would account for some of the variability in behavior; despite these differences, flies showed wing expansion in most trials. Because the main purpose of this experiment was to identify which population of neurons caused aggressive wing threat displays, we chose a simple metric (**Figure 3A**) – wingspan which we defined as the distance between the two wing tips – to analyze the effect of activating a given population of neurons. Activating all L320 neurons causes a large change in wingspan (**Figure 3B**) which is not observed in the control non-retinal flies (**Figure 3-S1B**). We also measured the effect of different light intensities and found that female flies appear more sensitive than male flies and will elicit robust wingspan increase at a light intensity (0.5 mW/cm^2^) at which males show only a weak and delayed wing threat (**Figure 3-S1C**). As the light intensity increases, latency to the initial wing response becomes faster (**Figure 3-S1D**), and the persistence of the wing response after the stimulus is turned off increases (**Figure 3-S1E**). A 1 mW/cm^2^ stimulus is sufficient to drive a robust wing response in both male and female flies with low latency and no change in baseline activity. When the stimulation intensity was increased to 2 mW/cm^2^, both male and female flies did not return to their initial completely closed-wing state. Rather, they will hold their wings slightly ajar for over 30 seconds (**Figure 3C and Figure 3-S1C/E**). Finally, after stimulation with a 4 mW/cm^2^ stimulus, the increase in wingspan becomes less robust and will sometimes not elicit wing movement among female flies (**Figure 3 S1C**). We utilized the 2 mW/cm^2^ stimulus to study the role of different neural populations because it generated the most robust response (**Figure 3-S1C**).

**Figure 3.**
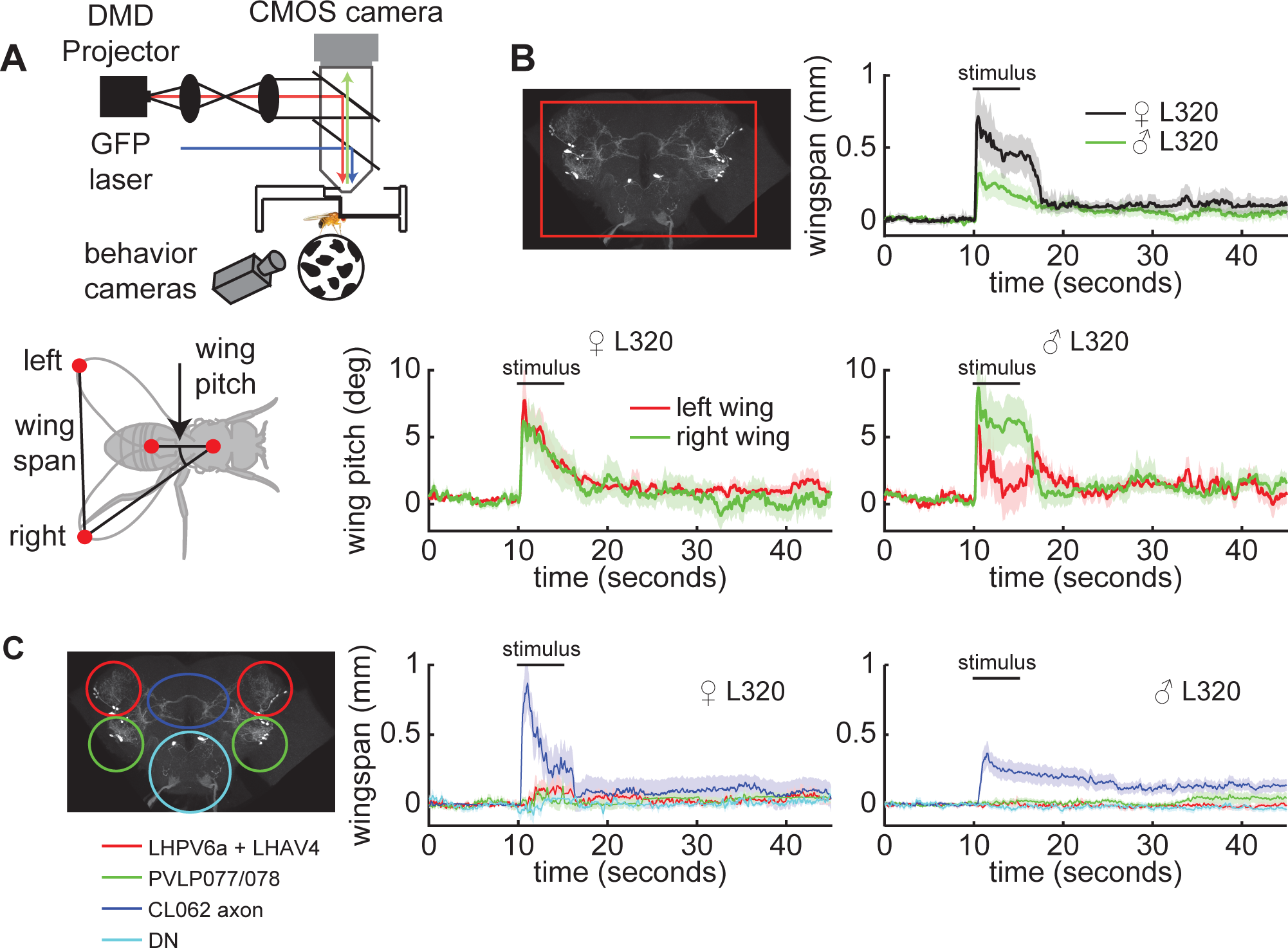
Optogenetic activation of a single population of neurons (CL062) drives wing threat in both females and males. **A.** Experimental setup. **Bottom:** Top down cartoon of a fly illustrating left/right wing pitch and wingspan. **B.** Optogenetic activation of all L320 neurons (L320 > UASChrimson) for males and females. (Females: n=11 trials across 5 flies, Males: n=26 trials across 7 flies). The wingspan (top right), and wing pitches following optogenetic stimulation using a 2 mW/cm² 617 nm light. **C.** Spatially targeted optogenetic activation of each type of neuron labeled by L320 > UAS Chrimson. Left: Illustration of regions targeted. Middle and right panels show baseline subtracted female and male wingspans respectively. (Females: n=12 trials across 5 flies, Males: n=29 trials across 7 flies). Neurons are activated using a 2 mW/cm² 617 nm light delivered at the neuron focal plane. Panels show mean +/− standard error of mean.

To assess which L320 neurons drive wing threat, we activated the LH, PVLP, CL062, and DN by changing the area over which the red light is turned on (**Figure 3C**). Since it was difficult to isolate the cell bodies of CL062 somas from the neurite of the LH and PVLP neurons within our preparation, we targeted the midline crossing portion of the CL062 axons (blue oval). We found that activation of the CL062 axons, but not the other neurons, drove a robust increase in wingspan (**Figure 3C and Figure 3-S2A**). The latency to the initial peak in wingspan and the wingspan habituation when only the CL062 axons are stimulated was most similar to that observed when performing brain-wide stimulation of all L320-split neurons at ∼ half the light intensity (**Figure 3-S2B/C**). Since channelrhodopsins, such as CsChrimson, can be expressed throughout the neuron and the midline crossing portion of the CL062 neurons that we targeted represents ∼half of the axon, this weaker behavioral response may be a consequence of activating only a subset of all Chrimson channels. Finally, we did observe a small increase in the wingspan in female flies when activating the LH and PVLP clusters of neurons (**Figure 3C**). This is likely due to off-target activation of CL062 somas since they are in close spatial proximity to the LH and PVLP neuron neurites within our experimental preparation (**Figure 3-S3B**).

To assess whether the behavior elicited by the L320 line in freely walking flies resulted from the activation of the CL062 neurons, we utilized a previously reported split-GAL4 line called Split^Thr^ (GMR22D03-AD ⋂ GMR20E08-DBD) that has been shown to drive male wing threats (Duistermars et al., 2018). We will refer to this line as CL062-split since it labels only the CL062 neurons and a small population of dorsal neurons that are not implicated in the behavior. Both female and male CL062-split flies performed the same set of actions; the time-course of the actions is similar to L320-split flies (**Figure 4 and Figure 4-S1A and SVideo 4**). During wing threat and wing extensions, flies raised their wings to qualitatively similar elevation and azimuth angles (**Figure 4-S1B**). Furthermore, when not performing a wing threat or extension, the CL062-split flies will also keep their wings slightly ajar (**Figure 4-S1C**). Despite the similarities, there are some differences in the temporal dynamics of the probability of observing lunges and the alert stance. Female CL062-split flies show a higher propensity to thrust immediately after stimulus onset while male CL062-split flies did not perform thrusts (**Figure 4**). A higher proportion of male CL062-split flies also stood in the alert stance throughout the stimulus period (**Figure 4B**). Female flies not fed with retinal displayed a muted amount of wing extension and alert stance, perhaps reflecting leaky optogenetic activation of neurons (**Figure 4-S2**). To confirm that the small increase in wingspan observed when we performed spatially restricted targeting of LH neurons was due to off-target activation of CL062 neurons and not the LH neurons themselves, we utilized two Janelia split-Gal4 lines called L188 (GMR47B03-AD ⋂ GMR30H02-DBD) and L2193 (VT060077-AD ⋂ VT029317-DBD) that label subsets of the LH neurons expressed by L320-split. L188 labels LHPV6a1 while L2193 labels LHPV6a3 neurons (Dolan et al., 2019). Optogenetic activation of these genetic lines did not result in wing threat, extension, lunging, or holding an alert stance (**Figure 4-S3**). It is important to note that activating LH neurons labeled by these lines did result in other behaviors. As these behaviors appear unrelated to aggression, we have not quantified them in this study. Taken together, these experiments suggest that the CL062 neurons are sufficient in driving wing threat, wing extension, thrusting, and alert stance in a temporally structured manner.

**Figure 4.**
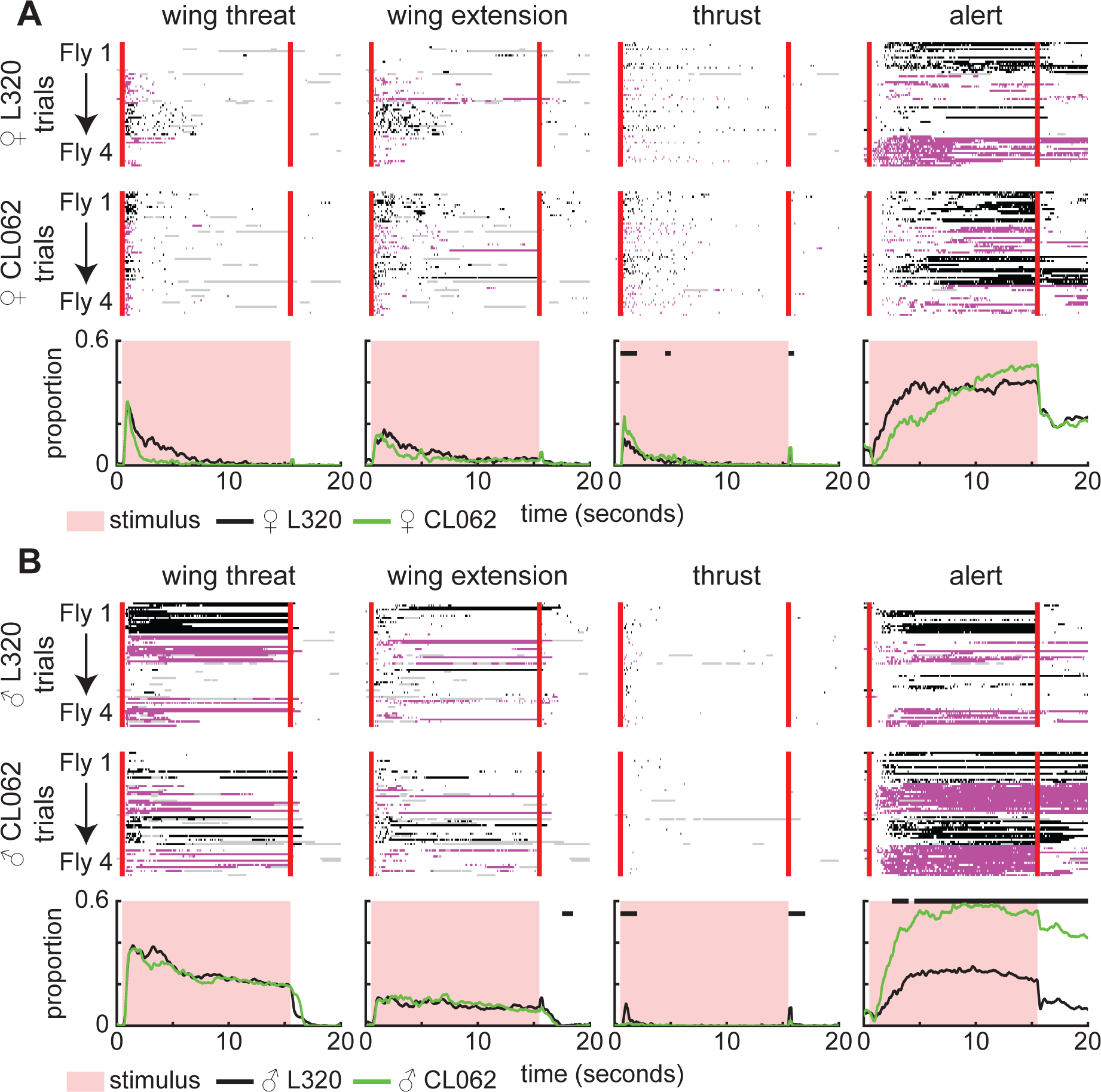
Activation of CL062 neurons using a more restrictive line reproduces L320 behavior. **A. Top rows:** Single trial ethogram of actions for 4 sample female L320 > UASChrimson and CL062-split > UASChrimson flies. Trials grouped by flies (fly 1 = trials 1-15, fly 2 = trials 16-30, etc). Red line indicates start and end of optogenetic stimulation period, which lasts for 15 seconds. Alternating black and purple blocks of trials represent different flies. Gray areas indicate frames where any of the tracked body parts used to identify the action had low tracking confidence (see methods). **Bottom row:** Proportion of trials where flies are performing each action. An 8.9 mW/cm² 617 nm light is turned on during the stimulus period. (Retinal fed: n=12 L320 flies, n=11 CL062 flies; 15 trials/fly). Black bars show time points where there is a significant difference between L320 > UASChrimson and CL062-split > UASChrimson flies (Wilcoxon rank sum test p<0.01, see methods). **B.** Same as **A**, but for male L320 > UASChrimson and CL062-split > UASChrimson flies. (Retinal fed: n=11 L320 flies, n=11 CL062 flies; 15 trials/fly).

#### Unilateral activation of CL062 drives bilateral wing threat followed by contralateral wing extension

The fact that these neurons can drive both independent movement of the wings as well as the extension of a single wing for long durations (**Figure 1-S3**) is surprising given that CL062 axons project to both hemispheres (**Figure 2**). To understand the source of wing extension, we performed unilateral activation of CL062 neurons in female flies (**Figure 5A and Figure 5-S1 and SVideo 5**). We found that unilateral stimulation drove an increase in ipsilateral wing pitch (**Figure 5B/C and Figure 5-S2A/B**). Surprisingly, like the ipsilateral wing, the contralateral wing also exhibited an initial increase in wing pitch that quickly adapted after 2 seconds. This response suggests a multi-timescale control of wings initiated by CL062 neurons: Initial activation of these neurons first drives a wing threat response utilizing both wings; the contralateral wing response habituates leaving only the ipsilateral wing extended; this long-lasting ipsilateral wing response is reminiscent of wing extension behavior observed during behavior in freely-walking flies. Interestingly, the behavioral response during bilateral activation of CL062 neurons is greater than the sum of unilateral activation, suggesting a potential form of non-linearity (**Figure 5D and Figure 5-S2C**).

**Figure 5.**
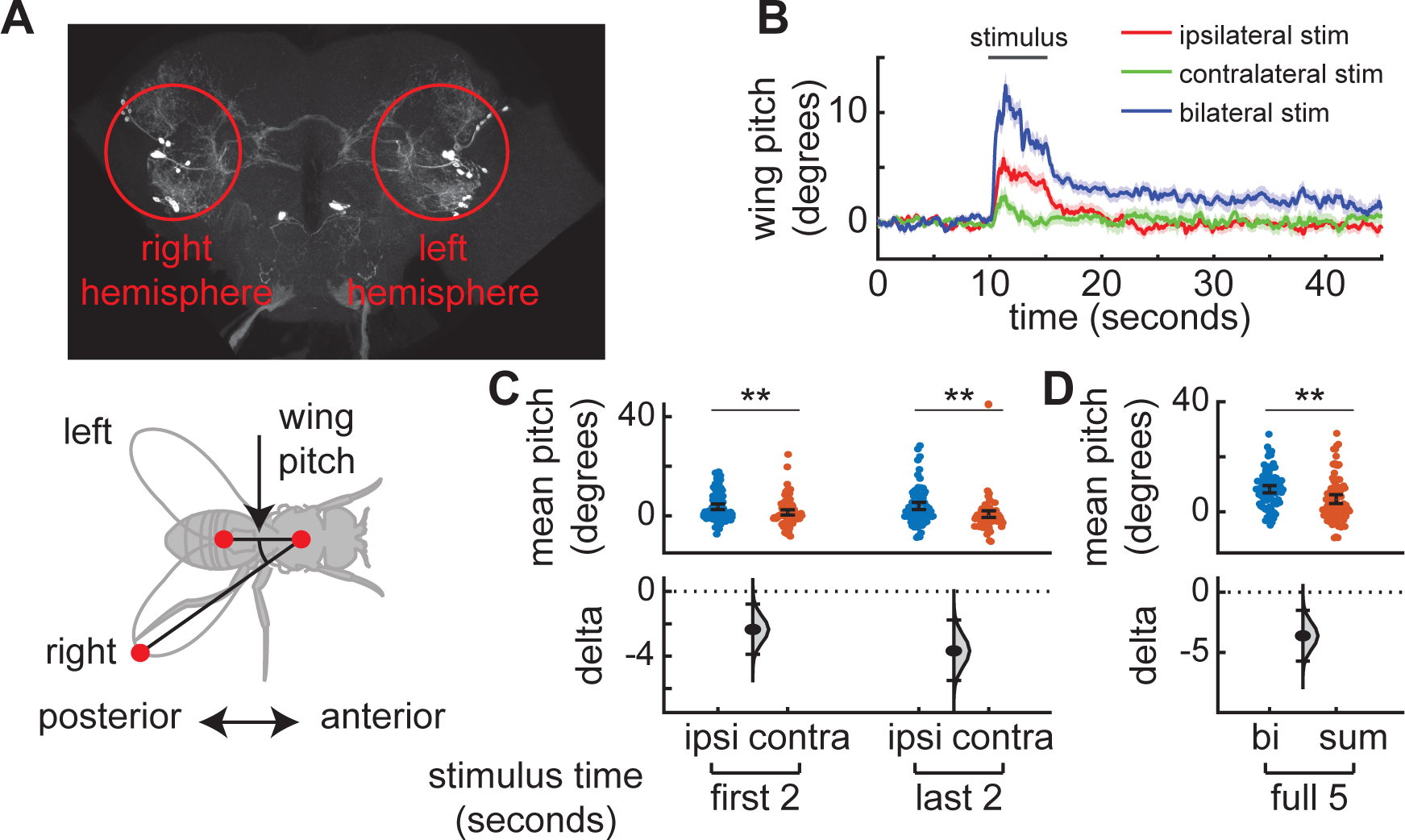
Unilateral activation of CL062 neurons shows that ipsilateral and contralateral wings threats are controlled differentially. **A. Top:** Experimental setup. Red circles show approximately the targeted regions during experimentation. Here, the anterior axis is facing out of the page. **Bottom:** Stick figures of a top down view of the fly showing the left and right wing pitch. **B.** Baseline subtracted wing pitch (both left and right) after ipsilateral, contralateral, and bilateral stimulation shows a fast increase in bilateral wing pitch followed by higher ipsilateral wing pitch. (Females: n=23 trials across 5 L320 > UASChrimson flies). Panel show mean +/− standard error of mean. Neurons are activated using a 2 mW/cm² 617 nm light delivered at the neuron focal plane. **C. Left:** Despite bilateral initial bilateral wing behavior on stimulus onset, the baseline subtracted wing pitch during the first 2 seconds of ipsilateral stimulus is higher than that of the contralateral stimulus. **Right:** The baseline subtracted wing pitch during the first 2 seconds of ipsilateral stimulus is higher than that of the contralateral stimulus. **D.** The baseline subtracted wing pitch during the entire 5 second bilateral stimulation period is significantly higher than the sum of the mean left and right pitch during the stimulus period(Wilcoxon rank sum test ** p<0.01).

#### CL062 likely drives downstream behaviors through multiple parallel descending neuron pathways

Since activation of CL062 neurons drives multiple actions, we next asked how the CL062 neurons are connected to downstream circuits to drive action. One possibility is that different CL062 neurons drive different subsets of actions in a modular manner. Action choice must be relayed from the brain to motor circuits in the ventral nerve cord through ∼ 1100 DNs (Hsu and Bhandawat, 2016). Therefore, if CL062 neurons drive actions in a modular manner, we would expect that different CL062 neurons will have stronger connections to different subsets of DNs. To examine whether there is modularity, we used Flywire and Natverse toolsets to obtain the connectivity of DNs that are postsynaptic of CL062 (Bates et al., 2020; Buhmann et al., 2021; Dorkenwald et al., 2022; Heinrich et al., 2018). These connections are based on a fully reconstructed female full adult fly brain (FAFB) and curated by the flywire community (Dorkenwald et al., 2022; Schlegel et al., 2023; Sven et al., 2023; Zheng et al., 2018). We found that each CL062 neuron makes at least 10 synapses with 3.7 DNs on average (**Figure 6A**). Although CL062 makes connections to non-descending DNs, the strong connections are more likely to be to DNs (**Figure 6B**) with these DNs making up ∼52% of total synaptic connections at this threshold. Only a single pair of DNs, DNpe050, is postsynaptic to every ipsilateral CL062 and most contralateral CL062 neurons (**Figure 6A**). This pair of neurons receives the largest output from the CL062 neurons when the outputs from all the CL062 neurons are summed together (**Figure 6B**). Besides this pair of DNs, the other DNs are postsynaptic to only small subsets of CL062 neurons within a single hemisphere; these DNs can be either ipsilateral or contralateral (**Figure 6A**). Finally, there is a single pair of DNs called pMP12 that putatively expresses *fru* according to the connectome data explorer Codex (Matsliah et al., 2023), a gene important for sexually dimorphic social behaviors (Gill, 1963; Vrontou et al., 2006). In a previous study, CL062 neurons have been shown to be connected to pMN1/DNp13, but these connections are through a smaller number of synapses and are not further considered in this study (Baker et al., 2022).

**Figure 6.**
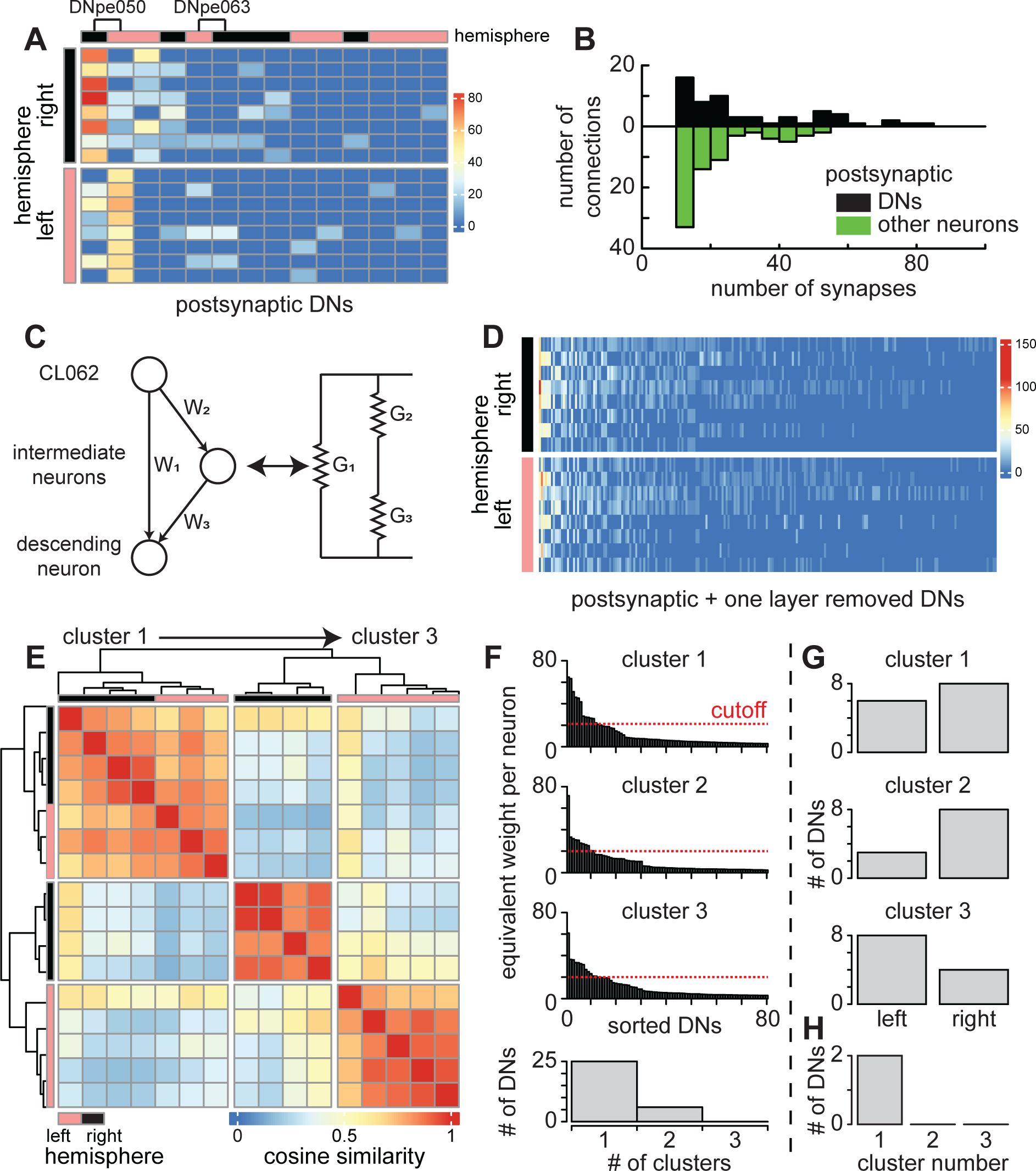
Modularity in CL062 connections to descending neurons (DNs). **A.** Number of synapses from each CL062 neuron to all direct postsynaptic DNs with at least 10 synapses. **B.** Number of synapses CL062 neurons make with DNs in comparison to all postsynaptic neurons. **C.** Schematic of equivalent neuron weights with up to a single intermediate neuron within a path. Single path weights is modeled as conductance of resistors in series. Equivalent weight is modeled as conductance of resistors in parallel. **D.** Equivalent weight from each CL062 neuron to all connected descending neurons. **E.** Heiarchical agglomative clustering of the cosine similarity between each pair of CL062 neuron to descending neuron connections. There are approximately three clusters of CL062 neurons. **F.** Sorted equivalent weights per neuron for each of the five clusters shows that each cluster connects strongly to a small number of descending neurons. A threshold of 20 was used to binarize strongly connected DNs. **Bottom:** Most DNs connect strongly to only a single cluster. **G.** The number of strongly connected DNs within each hemisphere to each cluster of CL062 neurons. **H.** Number of strongly connected DNs that putatively express fruitless for each cluster.

Despite being strongly connected to DNs, CL062 neurons also connect strongly to multiple interneurons, which may in turn connect to DNs (**Figure 6B**). These forms of indirect connections have been shown to be important for linking structure to functional correlation across the brain (Turner et al., 2021; Uzel et al., 2022). Therefore, we utilized a method based on resistor circuits to determine equivalent feedforward connection weights to all DNs with 0 to 1 intermediate neuron in between (**Figure 6C, methods**). There are two main features of this methodology. First, the equivalent weight is higher for DNs that can be reached via multiple intermediate neurons (i.e., parallel pathways). Second, the weight of a pathway is limited by the lowest edge weight between two neurons in the path. While CL062 can connect to a broader set of DNs through second-order connections, we found that these connections are still sparse and modular (**Figure 6D**). We quantified the similarity in CL062 neurons’ equivalent connections weights to DNs using cosine similarity and found that there are 3 clusters of single-hemispheric CL062 neurons (**Figure 6E**).

Since the clustering is based on cosine similarity, the presence of multiple clusters of CL062 neurons can be due to either distribution shape or modularity in DN connection identity. In the former case, a CL062 neuron that connects broadly to a distributed set of many DNs will have low cosine similarity to another CL062 neuron that connects specifically to a few DNs even if these DNs are strongly connected to both CL062 neurons. In the latter case, two CL062 neurons that have the same distribution of connections to different populations of DNs will have a low similarity. We found that CL062 neurons in each cluster make a similar exponentially decaying distribution of connections to different DNs. Using an equivalent weight threshold of 20 per CL062 neuron, we found that each cluster is strongly connected to less than 15 DNs (**Figure 6F**). Of these, most DNs make strong connections to only a single cluster of CL062 neurons and only three pairs of DNs make strong connections to two CL062 clusters (**Figure 6F and Figure 6-S1**). Furthermore, cluster 1, which is comprised of CL062 neurons from both hemispheres, is strongly connected to DNs in both hemispheres (**Figure 6G**). Meanwhile, clusters 2 and 3, which are comprised of single hemisphere CL062 neurons, are connected to more ipsilateral than contralateral neurons. Surprisingly, only cluster 1 CL062 neurons are strongly connected to the fruitless pMP12 (**Figure 6H and Figure 6-S1**).

The connectivity pattern that we observe here is consistent with a modular organization in the connections between CL062 neurons and descending neurons (DNs). We hypothesize that activating subsets of CL062 neurons will likely drive subsets of action. Since modular circuits often show mutual inhibition, we assessed the possibility of mutual inhibition between CL062 neurons as a mechanism for shaping the temporal progression of aggressive actions. Since CL062 neurons are all cholinergic, we again considered two layered connections between CL062 neurons. We found that CL062 neurons make sparse glutamatergic and GABAergic connections with each other (**Figure 6-S2**) consistent with weak mutual inhibition. Thus, the evidence for mutual inhibition is not strong; and the weak mutual inhibition is consistent with the idea that aggressive actions elicited by CL062 do not reflect strict progression and that multiple actions such as (wing threat and thrusts) can occur at the same time.

#### CL062 and aIPg connect to different sets of DNs and CL062 sparsely inhibits aIPg through the inhibition of pC1d

How are the CL062 neurons related to known aggression neurons? Till date, most neurons that mediate aggression are known to be part of the *fru+*/*dsx+* circuit. Are the CL062 neurons also part of the same circuit and either upstream or downstream to those neurons? It is also possible that aIPg/pC1 neurons are mutually antagonistic: During aggression, after approaching, female flies will transition into either a wing threat or a head butt. The choice of this action initiates two distinct action sequence loops (Nilsen et al., 2004). One action sequence loop involves wing threat and thrusting while the other involves head-butting and multiple types of fencing actions. It is possible that a mechanism for these two loops is mutual antagonism between CL062 which drives wing threat and aIPg that drives head-butting and fencing actions (Schretter et al., 2020).

In either case, it is likely that they also connect to different sets of DNs. Like CL062, aIPg neurons make connections to multiple DNs through one layer of intermediate neurons. If CL062 and aIPg are involved in circuits driving different actions during aggression, then we should find that they connect to different subsets of DNs. Indeed, when we compared the cosine similarity between all CL062 and aIPg neurons, we found that there is a low similarity between aIPg and CL062 neurons (**Figure 7A**). We next sought to determine the set of DNs that may be important for the separation of the action sequence loops by determining the set of DNs that contribute highly to segregating aIPg neurons from CL062 neurons. To accomplish this, we performed Principal Component Analysis (PCA) to find principal components (PCs) that explain most of the variance in the connections from aIPg and CL062 neurons to DNs. We found that the first 2 PCs partitioned the left and right-hemisphere aIPg and CL062 into different clusters (**Figure 7B**). We can construct a pair of near orthogonal support vectors in this space that defines a left/right axis and an aIPg/CL062 axis. The largest contributions of DNs in the direction of left and right hemisphere CL clusters are close to the left/right support axis (**Figure 7-S1**) reflecting that these DNpe050 neurons are the only pair of DNs that connect strongly to all ipsilateral and most contralateral CL062 neurons. Meanwhile, the largest loadings of DNs in the direction of the left and right hemisphere aIPg axis lie further away from the left/right support axis implying that these DNs are likely strongly connected with single hemisphere aIPg neurons.

**Figure 7.**
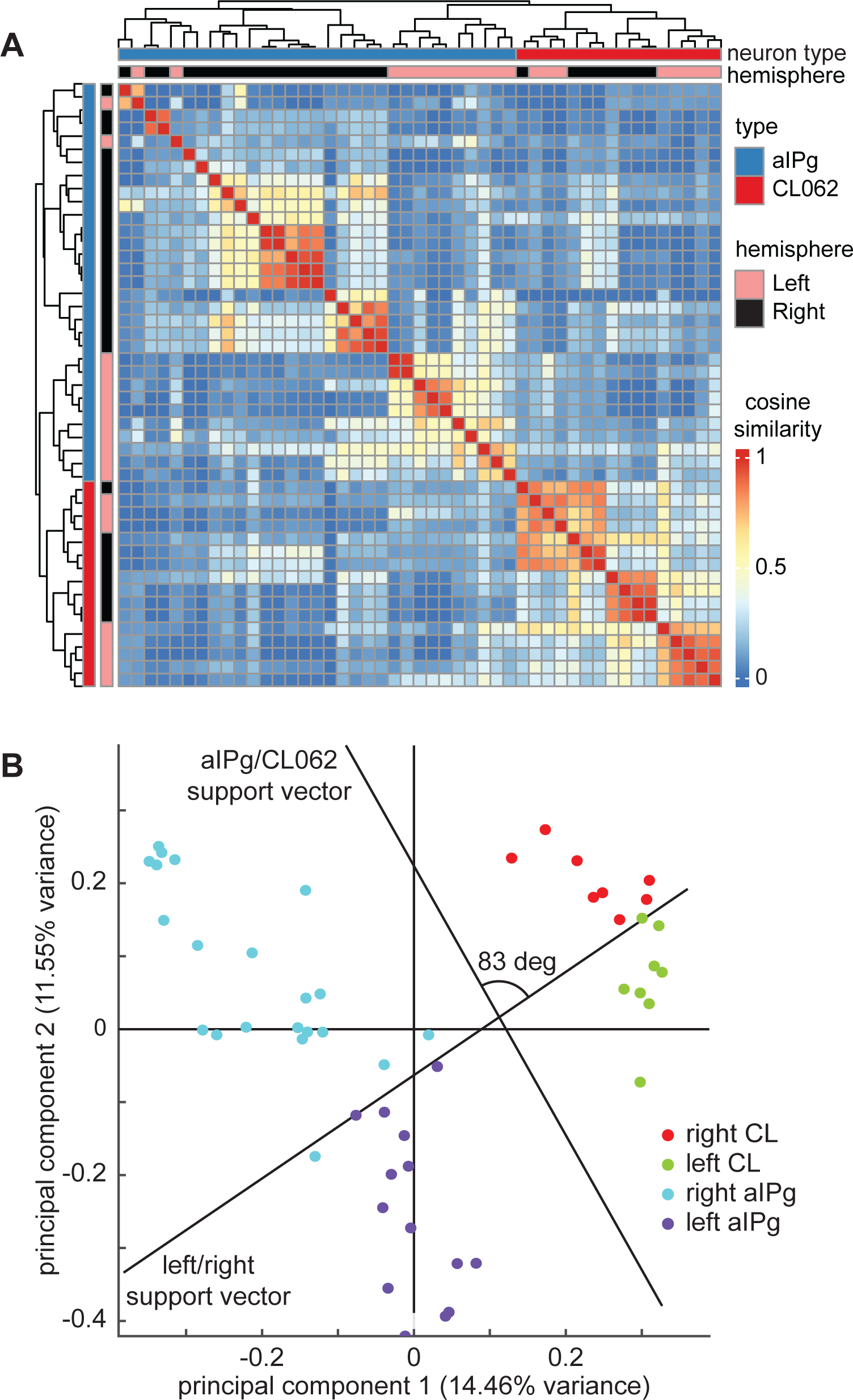
CL062 and aIPg likely drive actions through parallel descending neuron pathways. **A.** Hierarchical agglomative clustering of the cosine similarity between each pair of CL062 and aIPg neurons to descending neuron connections. CL062 do not make connections to similar descending neurons as aIPg. aIPg neurons that make a total equivalent weight of <20 to DNs were not considered. **B.** The first two principal components (PCs) of L2 normalized connections to descending neurons from CL and aIPg neurons. The first two PCs appear to separate left from right hemisphere neurons as well as aIPg from CL. The aIPg/CL062 and left/right support vectors had a 10-fold cross validation loss of 0 and 0.06 respectively. The two support vectors are nearly orthogonal.

Since CL062 and aIPg neurons drive different actions and connect strongly to different subsets of DNs, we next considered whether CL062 neurons act to inhibit neural circuits involving aIPg to drive the initial choice of wing threat over head-butting. A past study has found that aIPg receives strong inputs from a class of neurons called pC1d, which also drives a similar phenotype upon optogenetic activation (Deutsch et al., 2020; Palavicino-Maggio et al., 2019; Schretter et al., 2020). These pC1d neurons are part of and are recurrently connected to a group of 5 pC1 neurons per hemisphere that are involved in multiple social behaviors (Chiu et al., 2021; Koganezawa et al., 2016; Wang and Anderson, 2010; Zhou et al., 2014). Since CL062, aIPg, and pC1 neurons are all cholinergic, we again considered two-layer connections between these neuron types. We found that connections between individual CL062 neurons and aIPg/pC1 neurons were sparse (**Figure 8A**). CL062 neurons form a combination of GABAergic and Glutaminergic pathways to pC1d. In flies, GABA is the primary inhibitory neurotransmitter while glutamate has been shown in some visual and olfactory circuits to be inhibitory (Liu and Wilson, 2013; Molina-Obando et al., 2019). pC1d in turn excites two types of aIPg (aIPg1 and aIPg2) neurons as well as a subset of aIPg3 neurons. This connectivity pattern suggests a circuit motif where CL062 neurons could inhibit aIPg neurons indirectly through potential inhibition of pC1d. Meanwhile, aIPg neurons are sparsely connected to CL062 through GABAergic and Glutaminergic synapses. Interestingly, a subset of these aIPg3 neurons appears to receive input from aIPg2 neurons and make strong recurrent connections to each other that do not require other aIPg, pC1, or CL062 neurons. Finally, CL062 neurons do not appear to interact strongly with other classes of pC1 neurons even though pC1d appears to take strong feedforward input from pC1a and make strong recurrent pathways with pC1c and pC1e (**Figure 8**). In sum, although there are some interactions between these aggression circuits, they largely appear to be independent.

**Figure 8.**
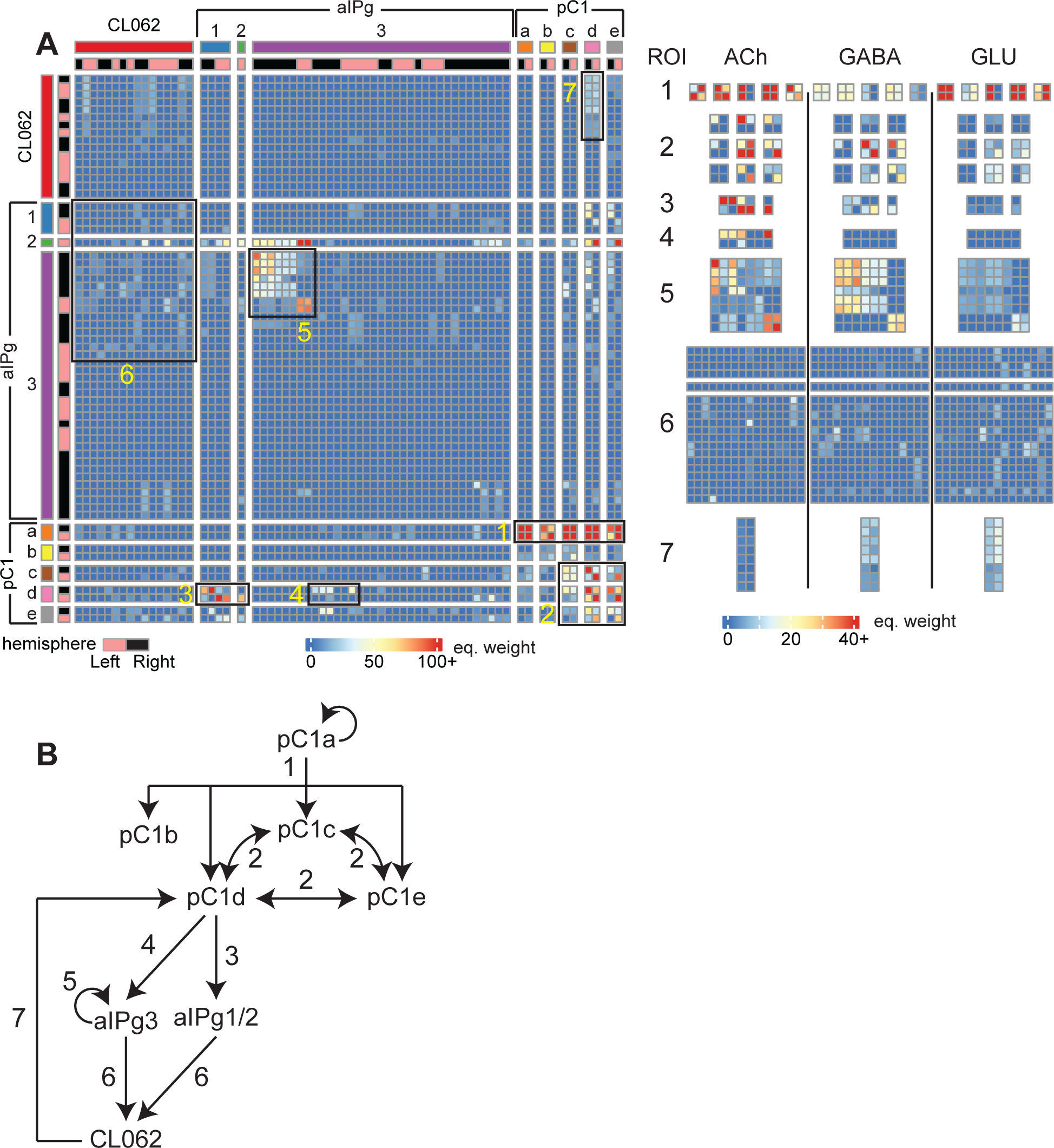
Potential pathways between CL062, aIPg, and pC1 neuron classes. **A. Left:** Weighted adjacency matrix (row -> column) showing equivalent weight between CL062, aIPg, and pC1 neurons. Seven regions are highlighed. **Right:** The equivalent weights for each of the seven regions broken down by neurotransmitter pathway. Since CL062, aIPg, and pC1 neurons are all cholinergic, GABA and Glutamate pathways arise from indirect pathways. **B.** Schematic showing the subset of potential pathways between neuron classes highlighted by the seven ROIs (numbers). Arrows indicate directionality. Pathways include both direct connections and one layer removed indirect connections via an intermediate neuron.

## Discussion

Several important findings emerge from our behavioral experiments and connectomics analysis. First, the CL062 neurons which were previously discovered to elicit aggressive actions in males (Duistermars et al., 2018) elicit aggressive actions in both males and females. Although activation of these neurons elicits the same action in both males and females, there are important differences between the male and female behaviors that mirror their aggressive actions in the presence of a target. Second, a target is not necessary for these actions. Third, the CL062 neurons do not express *fruitless* like the preponderance of other aggressive neurons (Duistermars et al., 2018). This lack of expression of genes that mediate much of the sexual dimorphism and the fact that these neurons are also not strongly connected to other known neurons that produce aggression implies that mechanisms underlying aggressive behavior in flies are more distributed than previously thought.

### Activating CL062 neurons in isolated freely-walking and head-fixed flies resemble aggressive actions observed in the presence of a mate

Are the actions that we observe here aggressive actions? Both the individual actions observed, and their time-course strongly suggest that the behavior observed here is aggression. At the level of individual aggressive actions, wing threats observed here, characterized by elevation of both wings above 45° are a hallmark of aggressive action (Nilsen et al., 2004), and distinct from wing extensions during courtship during which the wings move horizontally. Strikingly, although both males and females display wing threats upon activation of these neurons, the time course of the wing threats is sexually dimorphic: In males, wing threats last longer with some wing threat episodes lasting > 1 second (see **Figure 1-S5**). The same long-lasting wing threats have been observed by others (Duistermars et al., 2018; Nilsen et al., 2004). In contrast, female wing threats rarely lasted longer than 1 second. We observed this dimorphism not only in freely walking flies but also in head-fixed flies.

We also observe the many forms of thrusts reported by others: in some cases, thrusts simply involve a forward movement of the body without any concomitant movement of the legs, at other times; in another form of thrust, the forelegs are lifted, and the body elevates before snaping down; in yet another form, even the middle and back legs are lifted off the ground. These forms of thrusts are similar to those observed by others (Nilsen et al., 2004).

Apart from individual actions, the sequence of actions observed here has strong resemblance to the sequence observed during agonistic interactions between pairs of flies. Unlike the relative orderly progression during courtship, aggression involves a more complex structure of recurring behavioral sequences (Chen et al., 2002; Hoopfer, 2016; Nilsen et al., 2004). Within a single activation, a given action such as wing threat can occur by itself or along with multiple forms of thrust. Although a given action can occur throughout a trial, the probability of observing a given action changes over time; the change in probability is also sexually dimorphic: Males are more likely to show wing threat throughout the trial while females are more likely to show threats at the beginning of the trial. In contrast, females are more likely to thrust throughout the trial. These dimorphisms in behavior are reflective of dimorphism observed during natural agonistic interactions.

Taken together, these data suggest that CL062 neurons can not only mediate aggressive actions with a short latency but also accomplish this even in the absence of a target. Given that the individual actions resemble actions during natural aggressive behaviors and the sequence of actions resembles the sequence during natural behaviors, CL062 is likely a monomorphic node that orchestrates aggressive behaviors. The actions and their sequence appear to follow from the activation of this node.

### Implications for the organization of circuits that control aggressive behaviors in *Drosophila*

As aggressive behaviors are social behaviors, they are controlled by sexually dimorphic circuits. Many of the neurons involved in fly aggression are *fru*^+^/*dsx*^+^ and are important for mediating the choice between other social behaviors such as courtship. Since many social behaviors are dimorphic, these circuits are dimorphic as well. In this light, it is noteworthy that CL062 neurons are neither *fru*^+^ nor are they sexually dimorphic which makes understanding how the CL062 circuit relates to other aggression circuits critical. Since CL062 neurons drive aggressive behaviors in both males and females, there are implications for the circuit organization of aggressive behaviors in both.

Regarding the circuit organization of aggressive behaviors in the females, CL062 neurons appear to function independently of the previously characterized neurons for female aggression - aIPg and pC1d, and the relationship of CL062 to aggression has important differences from these neurons. One important difference is behavioral persistence. Activation of pC1d/e elicits persistent behavior in females(Chiu et al., 2023; Deutsch et al., 2020), in part, through its strong connections to aIPg. pC1d/e neurons also drive persistent activity in neurons expressing *Dsx* and *Fru* (Deutsch et al., 2020; Hui et al., 2023), and their activation produces minutes-long changes in behavioral state. In contrast, CL062 does not appear to have much long-lasting effect on aggression. Another important difference is that activation of aIPg and pC1d/e neurons in isolated flies has not been shown to drive aggressive behaviors (Deutsch et al., 2020; Palavicino-Maggio et al., 2019; Schretter et al., 2020); this lack of acute behavior is another fundamental difference from the CL062 neurons. One final difference is in the behaviors elicited by the two sets of neurons. The pC1d and/or aIPg neurons elicit a separate set of actions including headbutting, which do not occur concurrently with wing threat. One possibility is that CL062 and pC1d/aIPg might mediate the two parallel action sequence – one involving wing threat and the other involving headbutting(Nilsen et al., 2004). The presence of potentially inhibitory connections from CL062 to pC1d and from aIPg to CL062 provides a mechanism to inhibit the headbutt sequence loop once the decision for wing threat has been made (**Figure 8**). However, these connections are sparse; therefore, it is also possible that these two neuron groups represent independent aggression circuits that are recruited under different circumstances.

In males, we found that optogenetic activation of CL062 neurons drove actions in the absence of any sensory information. CL062 – referred to as AIP (anterior inferior protocerebrum) neurons in the other study - has already been implicated in male aggressive behaviors(Duistermars et al., 2018). In the other study, the authors did not observe lunging and concluded that these neurons mediate non-contact aggressive behaviors such as threats and not contact-behaviors such as lunges. This discrepancy could arise from several sources. First, there are differences in stimulation protocol between the two studies. Second, we performed our behaviors in isolated flies and not in the presence of a target. The presence of another fly might suppress contact aggressive actions. In a few pilot experiments that we performed with pairs of males; lunging in males was indeed suppressed; whenever we did observe lunges, they were not directed at the opponent. Third, it is possible that CL062 neurons are only important for lunging in the context of direct competition for a female during courtship. Consistent with this idea, CL062 neurons are responsive to the pulse component of the male courtship song(Baker et al., 2022). Pulse song can cause male wing threats(Sten et al., 2023). As in the case of females, CL062 neurons likely function in parallel to the well-studied aggression circuits in male - the sexually dimorphic neurons P1a neurons (Bath et al., 2014; Clemens et al., 2018; Hoopfer et al., 2015; Inagaki et al., 2014). The P1a neurons have been shown to drive immediate wing extension through direct activation of a pair of male-specific descending neurons (DNs) called pIP10 (Clemens et al., 2018; von Philipsborn et al., 2011) while activating a recurrent pathway that drives persistent lunging or courtship behaviors (depending on whether a male is present) for tens of minutes after the P1a neurons have stopped being active (Jung et al., 2020b). The P1a neurons are thought to be the switch in promoting an internal state of aggression or courtship due to this persistence of behavior. Given that P1 neurons are *fru*^+^ and some of their aggressive actions require male specific DNs, it is likely that they represent a circuit parallel to CL062 neurons. Another set of aggression producing neurons are the Tachykinin expressing aSP-g neurons that promote wing threat, lunging, and tussling toward other males (Asahina et al., 2014). As noted above, these neurons only make sparse connections to the CL062 neurons and are therefore also likely to function in parallel. In contrast, the tachykinin neurons work together with the P1a neuron to mediate aggression (Hoopfer et al., 2015).

### Hierarchical organization and parallel pathways of aggression

Work in neuroethology postulated hierarchically organized neural circuits underlying sexual behaviors (Dawkins, 1976; Tinbergen, 1951). At the top are “nervous centers” governing reproductive drive. These nervous centers then activate nervous centers governing the choice of competing behaviors such as aggression or courtship. As we descend the hierarchy, nervous centers represent increasingly specific patterns of movement (i.e., actions at level 3, then movement patterns within an action, then effectors, etc). Research in the last decade has provided evidence for circuits that underlie this hierarchical organization in multiple model systems. In mice, as postulated by neuroethologists, ventromedial hypothalamus contains dimorphic circuits that are implicated in both aggression and other sexual behaviors (Hashikawa et al., 2018; Lee et al., 2014; Lin et al., 2011; Pfaff and Sakuma, 1979a, b; Yang et al., 2013). In flies, too, the P1/pC1 cluster is sexually dimorphic and plays a large role in aggression, courtship, mate choice, and egglaying (Asahina, 2018; Chiu et al., 2021; Chiu et al., 2023; Deutsch et al., 2020; Hoopfer et al., 2015; Ishii et al., 2020; Koganezawa et al., 2016; Palavicino-Maggio et al., 2019; Rezával et al., 2016; Wang et al., 2021; Wang and Anderson, 2010; Wohl et al., 2020; Yang et al., 2019; Zhou et al., 2014). Recent work in flies has also shown that activity in these neurons can cause persistent changes in behavior (Chiu et al., 2023; Deutsch et al., 2020; Hoopfer et al., 2015; Jung et al., 2020b). Similar persistence has been observed in neurons in the ventromedial hypothalamus that drive multiple defensive behaviors (Kunwar et al., 2015; Wang et al., 2015). Where do the CL062 neurons fall within this hierarchy? Previously, it has been proposed that these CL062 neurons belong to a “level 3” nervous center since it was found that these neurons promoted non-contact behaviors – such as wing threats, charging, and turning – but not contact actions such as lunging in males (Duistermars et al., 2018). Connectomic results in this study, however, suggest that CL062 are likely to be independent: The set of descending neurons most likely to be activated by CL062 neurons appear to be different from those activated by the P1/pC1 cluster. This parallel circuit idea holds even when we trace the circuit through an intervening layer. Moreover, direct connections between P1/pC1 population and CL062 neurons are sparse suggesting that these circuits do not have strong interactions. It is still possible that CL062 are a part of the same hierarchy as the P1/pC1 population but they are not directly connected, or that the small number of direct connections have a disproportional effect.

Another possibility is that CL062 neurons mediate aggression that is not necessarily about competition over resources. As an example, females can occasionally move quickly towards a male with their wings extended as a nonreceptive response to male courtship(Sturtevant, 1915). Similarly, females have been shown to flick their wings and twist their abdomen to escape courting males(Manning, 1959). Thus, it is possible that CL062 neurons mediates other forms of aggression that require a rapid aggressive response rather than a long-lasting change in the state of the fly that characterizes aggression mediated by P1/pC1 neurons. In males, these neurons might play a role in competition during courtship. Consistent with this idea, CL062 neurons have recently been shown to be responsive to male courtship songs (Baker et al., 2022). The existence of an independent circuit for aggression makes sense, as aggression is not a “pure” dimorphic behavior in which males and females exhibit non-overlapping motor patterns to achieve a similar goal. Aggression is a “mixed” monomorphic-dimorphic behavior in which certain actions are shared and others dimorphic (Chiu et al., 2021). Recent studies have found a common circuit that mediates approach during aggression and recruits dimorphic circuits during the actual interaction phase of aggression (Chiu et al., 2021). However, many aggressive interactions are similar in males and females and might be mediated by independent circuits such as the CL062 neurons. If indeed CL062 neurons represent an independent aggression-promoting circuit, it is interesting to note that the downstream behavior still exhibits many characteristics of natural aggressive behaviors including sexual dimorphism. Regardless of whether CL062 is independent or not, it is noteworthy to find sexual dimorphism in a fly circuit that might not have a strong interaction with *fru*^+^ neurons.

## Acknowledgment

We would like to acknowledge the members of Bhandawat lab for discussions and for carefully reading the manuscript. This research was supported by RO1DC015827 (VB), RO1NS097881 (VB) and an NSF CAREER award (IOS-1652647 to VB), and an NIH F31NS120835-02 (LT).

We thank the Princeton FlyWire team and members of the Murthy and Seung labs, as well as members of the Allen Institute for Brain Science, for development and maintenance of FlyWire (supported by BRAIN Initiative grants MH117815 and NS126935 to Murthy and Seung). We also acknowledge members of the Princeton FlyWire team and the FlyWire consortium for neuron proofreading and annotation.

We thank the Drosophila Connectomics Group (PI G. Jefferis) for sharing their large scale proofreading and annotation in FAFB-FlyWire prior to publication. Proofreading and annotation in Cambridge were supported by Wellcome Trust Collaborative Awards (203261/Z/16/Z and 220343/Z/20/Z) to G. Jefferis; NIH BRAIN Initiative grant 1RF1MH120679-01 to D. Bock with G. Jefferis; and a Neuronex2 award to D. Bock and G. Jefferis (NSF 2014862, MRC MC_EX_MR/T046279/1). We specifically thank Varun Sane and Griffin Badalamente for proofreading and István Taisz and Dana Galili for their identification of aSP-g neurons.

## Author contributions

VB: Conceptualization; Supervision; Funding acquisition; Writing; analysis. LT: Conceptualization, Experimentation, analysis, Writing. VJB: Conceptualization, analysis, Writing; DA: Experiment and analysis.

## Competing interest statement

The authors declare that they have no competing interest or personal relationships that could have appeared to influence the work reported in this paper.

## METHODS

### CONTACT FOR REAGENT AND RESOURCE SHARING

Further information and requests for resources and reagents should be directed to and will be fulfilled by the lead contact, Dr. Vikas Bhandawat.

### DATA AND SOFTWARE AVAILABILITY

Data and analysis code supporting the study will be available after publication (as listed in the key resources table). Any additional information required to reanalyze the data is available from the corresponding author upon request.

### EXPERIMENTAL MODEL AND SUBJECT DETAILS

Flies were raised in 50 mL bottles of glucose media with ∼150 progeny/bottle (Archon Scientific D2). The number of progenies were controlled by the number of parent flies and the time parents were left on the food. A small sprinkle of active dry yeast was scattered on each bottle after removing the parents (1-3 days) to enrich the larvae diet. Bottles were placed in incubators set at 25°C (60% humidity) on a 12hr dark/ 12hr light cycle. Newly eclosed flies were raised in mixed 4 male + 4 female groups on 10 mL vials of D2 glucose media for control experiments. Retinal flies were raised with the same number of flies and put on food containing all-trans-retinal (0.02% by weight retinal) for optogenetic experiments. All vials were wrapped with aluminum foil to prevent retinal degradation and to keep conditions similar between retinal and control flies. Experiments were performed on flies 4 days after eclosion. Experimental flies were anesthetized on ice prior to placing them into the behavioral arenas.

### METHOD DETAILS

#### Key resources table

| REAGENT or RESOURCE | SOURCE | IDENTIFIER |
| --- | --- | --- |
| Chemicals, peptides, and recombinant proteins |  |  |
| All trans-Retinal | Sigma-Aldrich (MilliporeSigma) | Cat# R2500-1G |
| chicken anti-GFP | abcam | Cat# ab13970; RRID: AB_300798 |
| mouse anti-NC82 | DSHB | Cat# nc82; RRID: AB_2314866 |
| Alexa 488 anti-chicken | Thermo Fisher Scientific (Invitrogen) | Cat# A11039, RRID: AB_2534096 |
| Alexa 633 anti-mouse | Thermo Fisher Scientific (Invitrogen) | Cat# A21050, RRID: AB_2535718 |
| Dylight 550 anti-FLAG | Thermo Fisher Scientific (Invitrogen) | Cat# MA1-91878-D550; RRID: AB_1957945 |
| Dylight 488 anti-HA | Thermo Fisher Scientific (Invitrogen) | Cat# 26183-D488; RRID: AB_2533051 |
| Vectashield | Vector Laboratories | Cat# H-1000; RRID: AB_2336789 |
| Deposited data |  |  |
| Data (tracked behavior and connectomics) | This paper | To be decided during the publication process |
| Experimental models: Organisms/strains |  |  |
| *D. melanogaster*: w[1118]; P{y[+t7.7] w[+mC]=R33E02-p65.AD}attP40 | Bloomington Drosophila Stock Center | BDSC: 70189; FlyBase: FBst0070189 |
| *D. melanogaster*: w[1118]; P{y[+t7.7] w[+mC]=R47B03-GAL4.DBD}attP2 | Bloomington Drosophila Stock Center | BDSC: 68657; FlyBase: FBst0068657 |
| *D. melanogaster*: w[1118]; P{y[+t7.7] w[+mC]=R22D03-p65.AD}attP40 | Bloomington Drosophila Stock Center | BDSC: 69869; FlyBase: FBst0069869 |
| *D. melanogaster*: w[1118]; P{y[+t7.7] w[+mC]=R20E08-GAL4.DBD}attP2 | Bloomington Drosophila Stock Center | BDSC: 69771; FlyBase: FBst0026818 |
| *D. melanogaster*: w[1118]; P{y[+t7.7] w[+mC]=R47B03-p65.AD}attP40/CyO; P{y[+t7.7] w[+mC]=R30H02-GAL4.DBD}attP2/TM6B, Tb[1] | Bloomington Drosophila Stock Center | BDSC: 86709; FlyBase: FBst0086709 |
| *D. melanogaster*: w[1118]; P{y[+t7.7] w[+mC]=VT060077-p65.AD}attP40; P{y[+t7.7] w[+mC]=VT029317-GAL4.DBD}attP2/TM6B, Tb[1] | Bloomington Drosophila Stock Center | BDSC: 86634; FlyBase: FBst0086634 |
| *D. melanogaster*: w[1118] P{y[+t7.7] w[+mC]=20XUAS-IVS-CsChrimson.mVenus}attP18 | Bloomington Drosophila Stock Center | BDSC: 55134; FlyBase: FBst0055134 |
| *D. melanogaster*: w[1118] P{y[+t7.7] w[+mC]=hs-FLPG5.PEST}attP3; PBac{y[+mDint2] w[+mC]=10xUAS(FRT.stop)myr::smGdP-HA}VK00005 P{y[+t7.7] w[+mC]=10xUAS(FRT.stop)myr::smGdP-V5-THS-10xUAS(FRT.stop)myr::smGdP-FLAG}su(Hw)attP1 | Bloomington Drosophila Stock Center | BDSC: 64085; FlyBase: FBst0064085 |
| Software and algorithms |  |  |
| MATLAB r2022a | MathWorks | RRID: SCR_001622 |
| Python v3.8.17 | Anaconda (distributor) | RRID: SCR_008394 |
| RStudio (R v4.1.1) | Posit | RRID: SCR_000432 |
| DeepLabCut v2.3.5 | Mathis et al., 2018; Nath, Mathis et al., 2019; Lauer et al., 2022 | https://github.com/DeepLabCut/DeepLabCut RRID: SCR_021391 |
| Natverse and other connectome-related packages | Bates et al., 2020 | https://natverse.org/ |
| JAABA | Kabra, Robie, et al., 2012 | https://jaaba.sourceforge.net/ |
| Psychtoolbox 3.0 | Brainard, 1997; Pelli, 1997; Kleiner, Brainard, Pelli 2007 | http://psychtoolbox.org/ |
| DMD acquisition code and DLT triangulation GUI | This paper | https://github.com/bhandawatlab/DMDOptogeneticsExperiments |
| Analysis code and algorithms (behavior analysis, and connectomics) | This paper | https://github.com/bhandawatlab/2023_CL062_Tao_Bhandawat |

#### Freely-walking behavior (Figures 1 and 4)

##### Experimental setup

Behavioral experiments were performed in a setup similar to the one previously used in the lab (Chun et al., 2021). Flies were briefly immobilized using ice and placed onto a No 1 coverslip. A 20 x 10 x10 mm rectangular box made from cut microscope slides was secured over the fly using tape before the entire arena was held horizontally over a 45-degree mirror using clamps. The entire behavioral setup is enclosed in a Styrofoam box and a blackout curtain was draped over the box. The temperature surrounding the behavioral arena is maintained at 25 (+/−1 range) degrees Celsius using a heating blanket (Oven Industries 5R1-013). Flies were given a 30-minute acclimation period prior to the start of the experiment. We performed 15 trials per fly with a 5-minute rest period in between trials. Each trial lasted 30.5 seconds with a 617 nm red light (Thorlabs M617L3) triggered to turn on between the 0.5-second and 15.5-second mark. The light intensity was measured at 8.9 mW/cm^2^. Videos were captured at 100 Hz using a camera (Basler acA1920-150um) and focused with a 35mm lens (Edmund Optics 67-716). An infrared filter (Hoya IR76N) was placed at the end of the lens. The arena was lit using an 850 nm infrared light source (Thorlabs M850LP1).

##### Preprocessing of behavioral videos

The single-camera videos captured in the mirror chamber were split into a side view and a bottom view. For each view, we calculated the background by taking the mean of each pixel across a subsample of frames. Since there are trials where the fly stays in a single place, we then took the overall background as the mean of the backgrounds calculated across all trials for a given fly. After background subtraction, we calculated the log coefficient of variation (LCV) for each pixel:

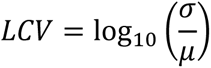

The foreground was defined as pixels with both an LCV of greater than 0.25 and a log standard deviation (*σ*) of less than 0.45. We then applied built-in MATLAB functions to perform contrast-limited adaptive histogram equalization on the foreground images followed by image sharpening (Zuiderveld, 1994). Finally, the noise pixels were added back to the processed foreground image to limit the loss of information from pixels containing parts of the fly that are improperly classified as noise.

##### Detection of body parts using DeepLabCut (DLC)

The head, thorax, abdomen, and left and right wingtips were detected using DLC (Mathis et al., 2018). We used the inbuilt k-means clustering to pick out unique frames and manually annotated the body parts in each frame. We trained a single Resnet-50 (He et al., 2016) for all corresponding cameras/views and both male and female flies. Prior to training, images were augmented using imgaug (Jung et al., 2020a). After each round of training, the network with the lowest mean average Euclidean error (MAE) in the testing dataset was applied to full videos and frames where tracked features had a confidence lower than 0.8 was extracted. We then used k-means clustering on these low-confidence frames to select the next round of refinement frames. The final network was trained with a 702 and 37 image training/testing set (95% training set) and had an MAE of 4.74 pixels (100.68 µm).

##### Triangulation

A micromanipulator was used to move a fluorescent microbead fixed at the end of a pulled glass micropipette. A custom MATLAB graphical user interface (GUI) was created to capture images from both the side and bottom view of the microbead at 72 positions in a box in XYZ space and to perform semiautomated detection of the microbeads. We performed triangulation using direct linear transformation (DLT). The DLT root mean squared error was 63.34 μm.

##### Definition of observables and actions

Using the tracked head, thorax, abdomen tip, and left and right wingtips, we calculated the fly’s speed, elevation angle, left wing pitch, and right-wing pitch. We calculated the speed as the displacement over time of the tracked thorax position. To calculate the left (and right) wing extension angle (*θ*), we first defined the body (head-thorax-abdomen) vector as the linear least squares vector in the direction from the abdomen to the head. The wing extension angle (*θ*) was then defined as the angle between the negative body vector 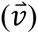 and the thorax wingtip vector 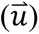:

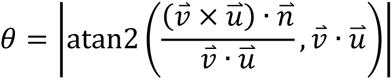

The elevation angle was defined as the angle between the body vector and the closest surface the fly is standing on. This angle is calculated using the same formulation as the wing extension angle. The closest surface that the fly is standing on is determined by first calculating the normal vector of each wall using the third column of the right singular matrix after performing singular value decomposition (SVD) on the wall plane centered at zero. Then, the distance to any given wall was calculated as the mean perpendicular distance between each endpoint (perpendicular projection of the head and abdomen positions onto the body axis) of the body axis and the wall.

To calculate the wing elevation angle, we first defined the plane formed by the tracked head, thorax, and abdomen point as the frontal plane with the normal vector pointing to the fly’s left side as 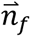. We can then project the wings and abdomen into the frontal plane:

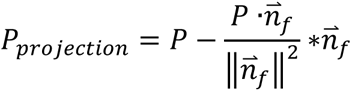

Where *P* is the wings or abdomen point and *P*_*projection*_ is the projection of that point on the plane. The wing elevation angles are then the angle between each wing projection to the abdomen projection in the frontal plane.

To calculate the wing azimuth angle, we defined the medial plane as the plane perpendicular to the frontal plane, coincident with the body vector, and with the normal vector 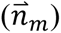 facing upwards. This was done by using the perpendicular projection of the head and abdomen positions onto the body axis, as well as another point defined by adding 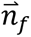 to the head projection on the body axis to define the plane. We can then project the wings and abdomen into this medial plane using the same formula as above but replacing 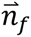 with 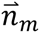. The wing azimuth angles are the angle between each wing projection to the abdomen projection in the medial plane.

Wing threat and extension were defined based on wing pitches (**Figure 1C and Figure 1-S1A/B**). First, a wing offset angle, which represents the extent that one wing is extended more than the other, is defined as the absolute angle between the line of unity (left pitch = right pitch) and the (x = left pitch, y = right pitch) vector. Wing threat was defined as when either the left- or right-wing pitch is greater than 45 degrees and the wing offset angle is less than 10 degrees. Wing extension was defined as when either wing has a pitch greater than 35 degrees and the wing offset angle is greater than 10 degrees. A third region of low-wing threat was defined, but not characterized in this study. This region is defined as either wing having a pitch greater than 35 degrees and less than 45 degrees and the wing offset was less than 10 degrees. The alert stance was defined by when the speed (thorax displacement) is less than 1 mm/s and the elevation angle was above 22.5 degrees. Thrusts were classified using a JAABA classifier with tracked observables (Kabra et al., 2013). We validated the performance of the classifier against manual annotations using 10-fold cross-validation and found 97.5% true positive and 96.9% true negative.

##### Analysis and statistical testing

Following DLC tracking, we marked tracked body parts with less than 70% confidence. Since two views were necessary for triangulation, the body parts were not triangulated for these instances and set to NaN. For continuous bouts of NaNs lasting less than 500 milliseconds (ms), we performed linear interpolation of the corresponding body part. When defining observables and actions, we set instances where dependent body parts were not triangulated or interpolated to NaN. For instance, if the side view of the left wing was marked as NaN at a particular time point, then the left-wing pitch, wing threat, and wing extension for that time point were set to NaN. These periods of NaNs are grayed out in the ethogram figures.

All ethograms as well as the observable traces (**Figure 1-S5**) were not smoothed. All statistical tests of actions over time were performed as follows. First, for each fly, we calculated the proportion of trials that the fly is performing the given action at each time point. This was smoothed by a 250 ms moving mean filter. To compare between two groups of flies, we looked at 500 ms non-overlapping time bins (i.e., 0-0.5 second, 0.5-1 second, etc.) and averaged each fly’s proportion over the time bin. We then calculated significance using a Wilcoxon rank sum test for each time bin using a significance cutoff of 0.01.

Latency to the first action bout was defined as the time from light onset to the start of the first bout of each action within a trial (**Figure 1-S6B**). The first bout was defined as the first bout that begins after the stimulus onset and lasts more than 30 ms.

During optogenetic stimulation, flies will keep their wings slightly ajar even when not performing wing threat or extension. As a first-order illustration of this, we fit Gaussian mixture models (GMM) to the wing pitch (**Figure 1-S1D and Figure 4-S1C**). We first fit a single Gaussian to the wing pitch in the 0.5 s period prior to light onset. We then fit a two-component GMM to 1.5 s (1 s overlapping) sliding window wing pitches. Time windows where the Bayesian information criterium of the two-component GMM is higher than that of a single Gaussian fit or where the difference in the mean of the two Gaussians in the two-component GMM fit was less than 5 degrees was fit using a single Gaussian. This analysis showed that:

1. There is a lower wing pitch distribution with an ever slightly higher mean than baseline.
2. Female flies showed faster habituation of wing pitch over the stimulus period.

#### DMD optogenetic experiments (Figures 3 and 5)

##### Experimental setup

Flies were anesthetized using ice before placing into a holder cut from a piece of aluminum foil. The head was stabilized using UV glue with the posterior brain facing upwards. After submersion in external saline (103 mmol/L NaCl, 5 mmol/L KCl, 5 mmol/L Tris, 10 mmol/L glucose, 26 mmol/L NaHC03, 1 mmol/L NaH2P04, 1.5 mmol/L CaCl2, 4 mmol/L MgCl2, osmolarity adjusted to 270–285 mOsm, bubbled with 95% O2/5% CO2 to pH 7.1–7.4), the cuticle was removed using forceps to expose the brain. A 20X/0.5 water immersion objective (Olympus UMPLFLN20XW) was used to visualize the brain. To activate different subpopulations of L320 neurons, we used a digital micromirror device (DMD) projector system (Wintech PRO 4500) to project stimulus patterns through the objective to the neurons of interest. A 700 mm projection lens (comes with the DMD) was placed at the output of the DMD. The light then passes through an achromatic doublet (Thorlabs AC254-035-A-ML) to correct for chromatic aberrations. Next, a neutral density filter (Thorlabs NE13A-A) was placed in the light path to lower the intensity of stimulation. The achromatic doublet and neutral density filter were attached to the 1X camera adaptor for a dual port (Olympus U-DP/U-DP1XC) enclosing a 50R/50T beam splitter (Edmund Optics 35-944). A schematic of the setup is shown in **Figure 3-S3A**.

Neurons were visualized using a CMOS camera (Hamamatsu Orco Flash 4.0). Prior to the start of the experiment, we took a z-stack of the fly brain to visualize the positions of neuron clusters expressing mVenus GFP (**Figure 3-S3B**). A custom GUI was made to define the circular stimulus ROIs (x, y, z position, and radius) and create a stimulus train using pre-calibrated light intensity curves for user-specified wavelengths. Red light (617 nm) was used for Chrimson activation. Psychtoolbox (Brainard and Vision, 1997) was hardware triggered using a national instrument data acquisition system (niDAQ) to present the stimulus train during experiments (NI USB-6363, 782258-01). The behavior was captured at 25Hz using 3 video cameras (2x Basler aCA800-510uc and 1x Basler acA1920-150um) and focused with 0.5X, 94 mm lens (Infinity Photo-Optical Company InfiniStix 194050). The fly was illuminated with two infrared 850 nm light sources (Lorex vq2121).

The transformation between the projector coordinates to CMOS camera pixel coordinates was calibrated using an 8×8 grid of evenly spaced white dots in the projector space. These dots were mapped to positions in the camera space using an affine transformation. The light intensity was calibrated by projecting a small circle in the center of the CMOS camera view with a stimulus train consisting of increasing normalized projector power output while a light meter (Thorlabs S121C and PM100USB) was placed underneath the objective to record the resulting light intensity. A No. 1 coverslip was placed on top of the light meter to immerse the 20X water immersion objective in external saline and the objective was focused using a checkerboard pattern. The light intensity curve for 617 nm is shown in **Figure 3-S3C**.

##### Detection of observables using DLC and triangulation

The head, thorax, first four strips on the abdomen, abdomen tip, and left and right wingtips in the 3 camera views were tracked using a single Resnet-50 model in DLC. The frame selection and training protocol was the same as that used for the freely walking DLC model. The final network was trained on 544 and 96 image training/testing sets (85% training set) and had an MAE of 4.03 pixels (38.69 µm). Calibration was performed for retinal-fed and non-retinal (control) fed experiments separately since the cameras were moved in between the days when the two sets of experiments were performed. We calibrated the DLT triangulation using 108 and 95 positions for the retinal-fed and control experiments respectively (we placed a microbead in 125 positions in a box in XYZ space, but not all positions were visible in at least 2 camera views). The DLT root mean squared error was 16.31 µm for retinal-fed flies and 48.53 µm for control flies.

##### Definition of observables and actions

While we tracked 9 body parts, only the thorax, first stripe, and left/right wingtips were necessary to calculate wing pitch angles and wingspan. The wing pitch angles were calculated in the same manner as the mirror chamber rig. The only difference is that the 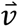 vector is defined as the vector from the thorax to the first stripe on the fly abdomen. The wingspan was calculated as the distance between the wingtips. The wing pitch angle(s) and wingspan were baseline subtracted by the mean value during the 10 seconds prior to each light stimulation period.

##### Analysis and statistical testing

The first peak in wingspan (**Figure 3-S1D**) was defined as the first peak in the baseline subtracted wingspan that is greater than 0.22 mm for males and 0.25 for females. A smaller threshold was chosen for males since female wings are longer and as a result, produce a larger wingspan. The ratio in the maximum male/female wingspan in the freely walking chamber was approximately 0.88.

**Figures 3-S2B/C, Figure 5C/D, and Figure 5-S2B/C** were analyzed using estimation methods to calculate mean, mean differences, and confidence intervals using a MatLab toolbox (Ho et al., 2019). Scatter plots show individual data points and corresponding error bars show mean and bootstrapped 95% confidence interval (resampled 10000 times, bias-corrected, and accelerated). 95% confidence interval for differences between means was calculated using the same bootstrapping methods. P-values were further generated using either Wilcoxon rank-sum or Wilcoxon sign rank tests.

#### Identifying neurons labeled by L320 (Figure 2)

##### Immunohistochemistry and imaging

Dissections and immunohistochemistry were carried out on Chrimson flies 3-5 days after eclosion. For Multicolor Flipout (MCFO) experiments, 3-day old flies were heat shocked at 37 degrees Celsius for 25-30 minutes. Dissections and immunohistochemistry were carried out 2-3 days after the heat shock. Flies were dissected in PBS, fixed in 2% paraformaldehyde for 55 minutes, blocked for 1.5 hours using 5% NGS, incubated in primary antibody in 5% NGS for at least overnight, incubated in secondary antibody in 5% NGS for at least 2 overnights and mounted using Vectashield (Vector Laboratories H-1000). After each step, the tissue was rinsed with PBS followed by 3 x 20-minute wash using 0.5 % Triton-X in PBS. Antibody incubation was performed on a 2D nutating shaker at 4 degrees Celsius. All other steps were performed on a 3D nutator at room temperature. The following primary antibodies were used: chicken anti-GFP (1:400 abcam ab13970), mouse anti-NC82 (1:20 DSHB). The following secondary antibodies were used: Alexa 488 anti-chicken (1:600 Invitrogen A11039), Alexa 633 anti-mouse (1:400 Invitrogen A21050). The following conjugate antibodies were used for MCFO experiments: Dylight 550 anti-FLAG (1:200 Invitrogen MA1-91878-D550), Dylight 488 anti-HA (1:200 Invitrogen 26183-D488). Fluorescence images were acquired using a Zeiss LSM700 inverted microscope confocal microscope.

#### Brain registration and neuron matching (Figure 2)

For the identification of L320 neuron IDs, we performed MCFO on female brains. We used CMTK to register dissected female brains to the JFRC2 template brain space. After registration to the JFRC2 template, we performed neuron tracing using the simple neurite tracer in ImageJ (Arshadi et al., 2021). The traced neurons were transformed to the hemibrain space (JRCFIB2018F) via JFRC2010 and JRC2018F using the neuroanatomy toolbox (NAT) (Bates et al., 2020; Bogovic et al., 2021). We then calculated a similarity score between transformed neurons to all hemibrain neurons using NBLAST (Costa et al., 2016). Hemibrain neurons with a NBLAST score of greater than 2500 were visually inspected to determine the most likely neuron match.

#### Connectomics analysis (Figures 6 and 7)

##### Choice of neuron IDs

We utilized Codex (Snapshot 630 April 2023 release) to obtain a list of neurons that are classified as CL062, pC1, aIPg, aSPg, and DNs (Matsliah et al., 2023). For CL062 neurons, we further stipulated that they crossed the midline and were given the community label of AIP since that was the given name to these neurons in the paper that first characterized the CL062-split genetic line (Duistermars et al., 2018). A list of neuron IDs with their corresponding type and hemisphere can be found in Table 1. This list does not include all DNs, only the ones mentioned by name or with morphology shown in the figures.

##### Determining equivalent connection weights

We utilized fafbseg to retrieve FAFB reconstructions from the production version of flywire (queried June 2023) to perform all connectomics analysis (Buhmann et al., 2021; Dorkenwald et al., 2022; Heinrich et al.; Li et al., 2020; Zheng et al., 2018). We set a cutoff of 10 synapses between any two neuron pairs for all connectivity analysis and considered the number of synapses between neurons as the edge weight between neurons. We utilized a resistor circuit approach to model equivalent (total) weight between two neurons that considers indirect paths through any arbitrary number of intermediate neurons. Under this formulation, the edge weights are treated as the conductance of a resistor. Therefore, the path weight for a single path between two neurons that passes through multiple intermediate neurons (series) is the inverse of the sum of the inverses of the individual weights between neurons in the path. Meanwhile, the equivalent weight for all paths between two neurons is simply the sum of all the individual path weights. In this paper, we considered only direct paths or paths with up to one intermediate neuron.

We used fafbseg to retrieve neurotransmitter/modulator (NT for brevity) predictions for each post-synaptic site from Flywire using a cleft score of 0 (Nils et al., 2023). These include GABA, acetylcholine, glutamate, octopamine, serotonin, and dopamine. The most confident NT was assigned to each synaptic site. The NT predictions between two neurons were defined as a 6-dimensional vector representing the number of synapses. Given a two layered path between two neurons (i.e., 3 nodes with 2 edges in series), there are 36 total permutations of path types (e.g., GABA->Ach, Ach->GABA, etc). However, since almost all synaptic assignments for CL062, aIPg, and pC1 neurons were cholinergic, in practice, the number of likely permutations was at most 6. Following our definition of equivalent weights, the equivalent weight associated with any permutation of NT i to NT j is:

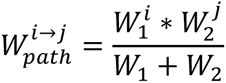

Where *W*_1_ and *W*_2_ are the total number of synapses (i.e., weight) associated with each edge of the path. 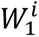 and 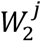 are the total number of synapses assigned to the i^th^ and j^th^ NT for the two path edges respectively.

##### Clustering/PCA/SVM analysis

CL062 were clustered by their connections to DNs by utilizing cosine similarity. Starting from the matrix defining the equivalent weight from CL062 (rows) to DNs (columns), cosine similarity is performed by performing a Euclidean normalization on each row. Next, the cosine similarity is defined as the cosine of the angle between each pair of rows. After performing cosine similarity, we performed hierarchical agglomerative clustering to cluster the Cl062 neurons into 3 clusters. For each cluster, we calculated the average equivalent weight for each CL062 neuron in the cluster to each DN (**Figure 6F**). A threshold of 20 was set because this was an approximate elbow point in the distribution of equivalent weights. The same process was repeated for building the similarity matrix for all alPg and CL062 neurons based on connections to DNs (**Figure 7A**).

Principal component analysis (PCA) was performed on Euclidean normalized equivalent weight matrix from CL062, aIPg, and pC1d/e to all descending neurons. The first two PCs explain 14.46 and 11.55 percent of the variance respectively. The PC scores were scaled by dividing by the maximum absolute value of the first 2 PC scores and then multiplying by the maximum distance of the loadings for the first 2 PCs. This scaling allows for viewing the loadings and the PC score in a biplot. Despite being unsupervised, the first two principal components (PCs) separated the left/right CL062 and aIPg neurons into different clusters. We created two support vector machines (SVM) in the PC space; one support vector to differentiate based on hemisphere and another to differentiate based on aIPg vs CL062. We used 10-fold cross validation to obtain a loss of 0.06 and 0 for the hemisphere and class support vectors respectively. Both PCA and SVM were performed using standard MATLAB function.

## Notes

### Competing Interest Statement

The authors have declared no competing interest.

## References

Anderson, D.J. (2016). Circuit modules linking internal states and social behaviour in flies and mice. Nat Rev Neurosci 17, 692–704.

Arshadi, C., Günther, U., Eddison, M., Harrington, K.I.S., and Ferreira, T.A. (2021). SNT: a unifying toolbox for quantification of neuronal anatomy. Nature methods 18, 374–377.

Asahina, K. (2017). Neuromodulation and strategic action choice in Drosophila aggression. Annual review of neuroscience 40, 51–75.

Asahina, K. (2018). Sex differences in Drosophila behavior: Qualitative and Quantitative Dimorphism. Curr Opin Physiol 6, 35–45.

Asahina, K., Watanabe, K., Duistermars, B.J., Hoopfer, E., González, C.R., Eyjólfsdóttir, E.A., Perona, P., and Anderson, D.J. (2014). Tachykinin-expressing neurons control male-specific aggressive arousal in Drosophila. Cell 156, 221–235.

Baker, C.A., McKellar, C., Pang, R., Nern, A., Dorkenwald, S., Pacheco, D.A., Eckstein, N., Funke, J., Dickson, B.J., and Murthy, M. (2022). Neural network organization for courtship-song feature detection in Drosophila. Current Biology 32, 3317–3333. e3317.

Bates, A.S., Manton, J.D., Jagannathan, S.R., Costa, M., Schlegel, P., Rohlfing, T., and Jefferis, G.S.X.E. (2020). The natverse, a versatile toolbox for combining and analysing neuroanatomical data. eLife 9, e53350.

Bath, D.E., Stowers, J.R., Hörmann, D., Poehlmann, A., Dickson, B.J., and Straw, A.D. (2014). FlyMAD: rapid thermogenetic control of neuronal activity in freely walking Drosophila. Nature Methods 11, 756–762.

Bogovic, J.A., Otsuna, H., Heinrich, L., Ito, M., Jeter, J., Meissner, G., Nern, A., Colonell, J., Malkesman, O., Ito, K., et al. (2021). An unbiased template of the Drosophila brain and ventral nerve cord. PLOS ONE 15, e0236495.

Brainard, D.H., and Vision, S. (1997). The psychophysics toolbox. Spatial vision 10, 433–436.

Buhmann, J., Sheridan, A., Malin-Mayor, C., Schlegel, P., Gerhard, S., Kazimiers, T., Krause, R., Nguyen, T.M., Heinrich, L., Lee, W.-C.A., et al. (2021). Automatic detection of synaptic partners in a whole-brain Drosophila electron microscopy data set. Nature Methods 18, 771–774.

Certel, S.J., and Kravitz, E.A. (2012). Scoring and analyzing aggression in Drosophila. Cold Spring Harb Protoc 2012, 319–325.

Chen, S., Lee, A.Y., Bowens, N.M., Huber, R., and Kravitz, E.A. (2002). Fighting fruit flies: A model system for the study of aggression. Proceedings of the National Academy of Sciences 99, 5664–5668.

Chiu, H., Hoopfer, E.D., Coughlan, M.L., Pavlou, H.J., Goodwin, S.F., and Anderson, D.J. (2021). A circuit logic for sexually shared and dimorphic aggressive behaviors in Drosophila. Cell 184, 507–520.e516.

Chiu, H., Robie, A.A., Branson, K.M., Vippa, T., Epstein, S., Rubin, G.M., Anderson, D.J., and Schretter, C.E. (2023). Cell type-specific contributions to a persistent aggressive internal state in female Drosophila (eLife Sciences Publications, Ltd).

Chun, C., Biswas, T., and Bhandawat, V. (2021). Drosophila uses a tripod gait across all walking speeds, and the geometry of the tripod is important for speed control. eLife 10, e65878.

Clemens, J., Coen, P., Roemschied, F.A., Pereira, T.D., Mazumder, D., Aldarondo, D.E., Pacheco, D.A., and Murthy, M. (2018). Discovery of a New Song Mode in Drosophila Reveals Hidden Structure in the Sensory and Neural Drivers of Behavior. Curr Biol 28, 2400–2412.e2406.

Costa, M., Manton, J.D., Ostrovsky, A.D., Prohaska, S., and Jefferis, G.S. (2016). NBLAST: Rapid, Sensitive Comparison of Neuronal Structure and Construction of Neuron Family Databases. Neuron 91, 293–311.

Dankert, H., Wang, L., Hoopfer, E.D., Anderson, D.J., and Perona, P. (2009). Automated monitoring and analysis of social behavior in Drosophila. Nature Methods 6, 297–303.

Darwin, C. (1874). The descent of man, and selection in relation to sex, Vol 1 (Murray).

Dawkins, R. (1976). Hierarchical organisation: A candidate principle for ethology. In Growing points in ethology (Oxford, England: Cambridge U Press).

Deutsch, D., Pacheco, D., Encarnacion-Rivera, L., Pereira, T., Fathy, R., Clemens, J., Girardin, C., Calhoun, A., Ireland, E., Burke, A., et al. (2020). The neural basis for a persistent internal state in Drosophila females. eLife 9, e59502.

Dolan, M.-J., Frechter, S., Bates, A.S., Dan, C., Huoviala, P., Roberts, R.J.V., Schlegel, P., Dhawan, S., Tabano, R., Dionne, H., et al. (2019). Neurogenetic dissection of the Drosophila lateral horn reveals major outputs, diverse behavioural functions, and interactions with the mushroom body. eLife 8, e43079.

Dorkenwald, S., McKellar, C.E., Macrina, T., Kemnitz, N., Lee, K., Lu, R., Wu, J., Popovych, S., Mitchell, E., Nehoran, B., et al. (2022). FlyWire: online community for whole-brain connectomics. Nature Methods 19, 119–128.

Duistermars, B.J., Pfeiffer, B.D., Hoopfer, E.D., and Anderson, D.J. (2018). A Brain Module for Scalable Control of Complex, Multi-motor Threat Displays. Neuron 100, 1474–1490.e1474.

Dulac, C., and Kimchi, T. (2007). Neural mechanisms underlying sex-specific behaviors in vertebrates. Curr Opin Neurobiol 17, 675–683.

Gill, K.S. (1963). A mutation causing abnormal courtship and mating behavior in males of Drosophila melanogaster. Am Zool 3, 507.

Hashikawa, K., Hashikawa, Y., Lischinsky, J., and Lin, D. (2018). The Neural Mechanisms of Sexually Dimorphic Aggressive Behaviors. Trends Genet 34, 755–776.

He, K., Zhang, X., Ren, S., and Sun, J. (2016). Deep residual learning for image recognition. Paper presented at: Proceedings of the IEEE conference on computer vision and pattern recognition.

Heinrich, L., Funke, J., Pape, C., Nunez-Iglesias, J., and Saalfeld, S. Synaptic cleft segmentation in non-isotropic volume electron microscopy of the complete drosophila brain (Springer).

Heinrich, L., Funke, J., Pape, C., Nunez-Iglesias, J., and Saalfeld, S. (2018). Synaptic Cleft Segmentation in Non-Isotropic Volume Electron Microscopy of the Complete Drosophila Brain. Paper presented at: International Conference on Medical Image Computing and Computer-Assisted Intervention.

Ho, J., Tumkaya, T., Aryal, S., Choi, H., and Claridge-Chang, A. (2019). Moving beyond P values: data analysis with estimation graphics. Nature Methods 16, 565–566.

Hoffmann, A.A. (1987). A laboratory study of male territoriality in the sibling species Drosophila melanogaster and D. simulans. Animal Behaviour 35, 807–818.

Hoopfer, E.D. (2016). Neural control of aggression in Drosophila. Current Opinion in Neurobiology 38, 109–118.

Hoopfer, E.D., Jung, Y., Inagaki, H.K., Rubin, G.M., and Anderson, D.J. (2015). P1 interneurons promote a persistent internal state that enhances inter-male aggression in Drosophila. eLife 4, e11346.

Hsu, C.T., and Bhandawat, V. (2016). Organization of descending neurons in Drosophila melanogaster. Scientific Reports 6, 20259.

Hui, C., Alice, A.R., Kristin, M.B., Tanvi, V., Samantha, E., Gerald, M.R., David, J.A., and Catherine, E.S. (2023). Cell type-specific contributions to a persistent aggressive internal state in female *Drosophila*. bioRxiv, 2023.2006.2007.543722.

Inagaki, H.K., Jung, Y., Hoopfer, E.D., Wong, A.M., Mishra, N., Lin, J.Y., Tsien, R.Y., and Anderson, D.J. (2014). Optogenetic control of Drosophila using a red-shifted channelrhodopsin reveals experience-dependent influences on courtship. Nature Methods 11, 325–332.

Ishii, K., Wohl, M., DeSouza, A., and Asahina, K. (2020). Sex-determining genes distinctly regulate courtship capability and target preference via sexually dimorphic neurons. Elife 9.

Jung, A.B., Wada, K., Crall, J., Tanaka, S., Graving, J., Reinders, C., Yadav, S., Banerjee, J., Vecsei, G., and Kraft, A. (2020a). imgaug. GitHub: San Francisco, CA, USA.

Jung, Y., Kennedy, A., Chiu, H., Mohammad, F., Claridge-Chang, A., and Anderson, D.J. (2020b). Neurons that Function within an Integrator to Promote a Persistent Behavioral State in Drosophila. Neuron 105, 322–333.e325.

Kabra, M., Robie, A.A., Rivera-Alba, M., Branson, S., and Branson, K. (2013). JAABA: interactive machine learning for automatic annotation of animal behavior. Nature Methods 10, 64–67.

Klapoetke, N.C., Murata, Y., Kim, S.S., Pulver, S.R., Birdsey-Benson, A., Cho, Y.K., Morimoto, T.K., Chuong, A.S., Carpenter, E.J., Tian, Z., et al. (2014). Independent optical excitation of distinct neural populations. Nat Methods 11, 338–346.

Koganezawa, M., Kimura, K., and Yamamoto, D. (2016). The Neural Circuitry that Functions as a Switch for Courtship versus Aggression in Drosophila Males. Curr Biol 26, 1395–1403.

Kravitz, E.A., and Fernandez, M.P. (2015). Aggression in Drosophila. Behav Neurosci 129, 549–563.

Kunwar, P.S., Zelikowsky, M., Remedios, R., Cai, H., Yilmaz, M., Meister, M., and Anderson, D.J. (2015). Ventromedial hypothalamic neurons control a defensive emotion state. Elife 4.

Kurtovic, A., Widmer, A., and Dickson, B.J. (2007). A single class of olfactory neurons mediates behavioural responses to a Drosophila sex pheromone. Nature 446, 542–546.

Lee, H., Kim, D.-W., Remedios, R., Anthony, T.E., Chang, A., Madisen, L., Zeng, H., and Anderson, D.J. (2014). Scalable control of mounting and attack by Esr1+ neurons in the ventromedial hypothalamus. Nature 509, 627–632.

Li, P.H., Lindsey, L.F., Januszewski, M., Zheng, Z., Bates, A.S., Taisz, I., Tyka, M., Nichols, M., Li, F., Perlman, E., et al. (2020). Automated Reconstruction of a Serial-Section EM *Drosophila* Brain with Flood-Filling Networks and Local Realignment. bioRxiv, 605634.

Lin, D., Boyle, M.P., Dollar, P., Lee, H., Lein, E.S., Perona, P., and Anderson, D.J. (2011). Functional identification of an aggression locus in the mouse hypothalamus. Nature 470, 221–226.

Lischinsky, J.E., and Lin, D. (2020). Neural mechanisms of aggression across species. Nat Neurosci 23, 1317–1328.

Liu, W., Liang, X., Gong, J., Yang, Z., Zhang, Y.H., Zhang, J.X., and Rao, Y. (2011). Social regulation of aggression by pheromonal activation of Or65a olfactory neurons in Drosophila. Nat Neurosci 14, 896–902.

Liu, W.W., and Wilson, R.I. (2013). Glutamate is an inhibitory neurotransmitter in the Drosophila olfactory system. Proc Natl Acad Sci U S A 110, 10294–10299.

Lorenz, K. (1963). On aggression (Harcourt, Brace and World: New York).

Manning, A. (1959). The Sexual Behaviour of Two Sibling Drosophila Species. Behaviour, 123–145.

Manoli, D.S., Fan, P., Fraser, E.J., and Shah, N.M. (2013). Neural control of sexually dimorphic behaviors. Curr Opin Neurobiol 23, 330–338.

Mathis, A., Mamidanna, P., Cury, K.M., Abe, T., Murthy, V.N., Mathis, M.W., and Bethge, M. (2018). DeepLabCut: markerless pose estimation of user-defined body parts with deep learning. Nature Neuroscience 21, 1281–1289.

Matsliah, A., Sterling, A., Dorkenwald, S., Kuehner, K., Morey, R., Seung, H., and Murthy, M. (2023). Codex: Connectome Data Explorer.

Molina-Obando, S., Vargas-Fique, J.F., Henning, M., Gür, B., Schladt, T.M., Akhtar, J., Berger, T.K., and Silies, M. (2019). ON selectivity in the Drosophila visual system is a multisynaptic process involving both glutamatergic and GABAergic inhibition. eLife 8, e49373.

Nelson, R.J., and Trainor, B.C. (2007). Neural mechanisms of aggression. Nature Reviews Neuroscience 8, 536–546.

Nern, A., Pfeiffer, B.D., and Rubin, G.M. (2015). Optimized tools for multicolor stochastic labeling reveal diverse stereotyped cell arrangements in the fly visual system. Proceedings of the National Academy of Sciences 112, E2967–E2976.

Nils, E., Alexander Shakeel, B., Andrew, C., Michelle, D., Yijie, Y., Philipp, S., Alicia Kun-Yang, L., Thomson, R., Samantha, F.-M., Tyler, P., et al. (2023). Neurotransmitter Classification from Electron Microscopy Images at Synaptic Sites in Drosophila Melanogaster. bioRxiv, 2020.2006.2012.148775.

Nilsen, S.P., Chan, Y.-B., Huber, R., and Kravitz, E.A. (2004). Gender-selective patterns of aggressive behavior in Drosophila melanogaster. Proceedings of the National Academy of Sciences 101, 12342–12347.

Palavicino-Maggio, C.B., Chan, Y.-B., McKellar, C., and Kravitz, E.A. (2019). A small number of cholinergic neurons mediate hyperaggression in female Drosophila. Proceedings of the National Academy of Sciences 116, 17029–17038.

Pandolfi, M., Scaia, M.F., and Fernandez, M.P. (2021). Sexual Dimorphism in Aggression: Sex-Specific Fighting Strategies Across Species. Front Behav Neurosci 15, 659615.

Pfaff, D.W., and Sakuma, Y. (1979a). Deficit in the lordosis reflex of female rats caused by lesions in the ventromedial nucleus of the hypothalamus. J Physiol 288, 203–210.

Pfaff, D.W., and Sakuma, Y. (1979b). Facilitation of the lordosis reflex of female rats from the ventromedial nucleus of the hypothalamus. J Physiol 288, 189–202.

Rezával, C., Pattnaik, S., Pavlou, H.J., Nojima, T., Brüggemeier, B., D’Souza, L.A.D., Dweck, H.K.M., and Goodwin, S.F. (2016). Activation of Latent Courtship Circuitry in the Brain of Drosophila Females Induces Male-like Behaviors. Curr Biol 26, 2508–2515.

Scheffer, L.K., Xu, C.S., Januszewski, M., Lu, Z., Takemura, S.Y., Hayworth, K.J., Huang, G.B., Shinomiya, K., Maitlin-Shepard, J., Berg, S., et al. (2020). A connectome and analysis of the adult Drosophila central brain. Elife 9.

Schlegel, P., Yin, Y., Bates, A.S., Dorkenwald, S., Eichler, K., Brooks, P., Han, D.S., Gkantia, M., Dos Santos, M., Munnelly, E.J., et al. (2023). Whole-brain annotation and multi-connectome cell typing quantifies circuit stereotypy in Drosophila. bioRxiv.

Schretter, C.E., Aso, Y., Robie, A.A., Dreher, M., Dolan, M.-J., Chen, N., Ito, M., Yang, T., Parekh, R., Branson, K.M., et al. (2020). Cell types and neuronal circuitry underlying female aggression in Drosophila. eLife 9, e58942.

Scott, J.P. (1966). Agonistic Behavior of Mice and Rats: A Review. American Zoologist 6, 683–701.

Siwicki, K.K., and Kravitz, E.A. (2009). Fruitless, doublesex and the genetics of social behavior in Drosophila melanogaster. Current Opinion in Neurobiology 19, 200–206.

Sten, T.H., Li, R., Hollunder, F., Eleazer, S., and Ruta, V. (2023). Male-male interactions shape mate selection in *Drosophila*. bioRxiv, 2023.2011.2003.565582.

Sturtevant, A.H. (1915). Experiments on sex recognition and the problem of sexual selection in Drosoophilia. Journal of Animal Behavior 5, 351.

Sven, D., Arie, M., Amy, R.S., Philipp, S., Szi-chieh, Y., Claire, E.M., Albert, L., Marta, C., Katharina, E., Yijie, Y., et al. (2023). Neuronal wiring diagram of an adult brain. bioRxiv, 2023.2006.2027.546656.

Taisz, I., Donà, E., Münch, D., Bailey, S.N., Morris, B.J., Meechan, K.I., Stevens, K.M., Varela-Martínez, I., Gkantia, M., Schlegel, P., et al. (2023). Generating parallel representations of position and identity in the olfactory system. Cell 186, 2556–2573.e2522.

Tinbergen, N. (1951). The study of instinct (New York, NY, US: Clarendon Press/Oxford University Press).

Turner, M.H., Mann, K., and Clandinin, T.R. (2021). The connectome predicts resting-state functional connectivity across the Drosophila brain. Current Biology 31, 2386–2394.e2383.

Uzel, K., Kato, S., and Zimmer, M. (2022). A set of hub neurons and non-local connectivity features support global brain dynamics in C. elegans. Current Biology 32, 3443–3459.e3448.

von Philipsborn, A.C., Liu, T., Yu, J.Y., Masser, C., Bidaye, S.S., and Dickson, B.J. (2011). Neuronal Control of Drosophila Courtship Song. Neuron 69, 509–522.

Vrontou, E., Nilsen, S.P., Demir, E., Kravitz, E.A., and Dickson, B.J. (2006). fruitless regulates aggression and dominance in Drosophila. Nature Neuroscience 9, 1469–1471.

Wang, K., Wang, F., Forknall, N., Yang, T., Patrick, C., Parekh, R., and Dickson, B.J. (2021). Neural circuit mechanisms of sexual receptivity in Drosophila females. Nature 589, 577–581.

Wang, L., and Anderson, D.J. (2010). Identification of an aggression-promoting pheromone and its receptor neurons in Drosophila. Nature 463, 227–231.

Wang, L., Chen, I.Z., and Lin, D. (2015). Collateral pathways from the ventromedial hypothalamus mediate defensive behaviors. Neuron 85, 1344–1358.

Watanabe, K., Chiu, H., Pfeiffer, B.D., Wong, A.M., Hoopfer, E.D., Rubin, G.M., and Anderson, D.J. (2017). A Circuit Node that Integrates Convergent Input from Neuromodulatory and Social Behavior-Promoting Neurons to Control Aggression in Drosophila. Neuron 95, 1112–1128.e1117.

Wohl, M., Ishii, K., and Asahina, K. (2020). Layered roles of fruitless isoforms in specification and function of male aggression-promoting neurons in Drosophila. Elife 9.

Wu, F., Deng, B., Xiao, N., Wang, T., Li, Y., Wang, R., Shi, K., Luo, D.-G., Rao, Y., and Zhou, C. (2020). A neuropeptide regulates fighting behavior in Drosophila melanogaster. eLife 9, e54229.

Yamamoto, D., and Koganezawa, M. (2013). Genes and circuits of courtship behaviour in Drosophila males. Nat Rev Neurosci 14, 681–692.

Yang, C.F., Chiang, M.C., Gray, D.C., Prabhakaran, M., Alvarado, M., Juntti, S.A., Unger, E.K., Wells, J.A., and Shah, N.M. (2013). Sexually dimorphic neurons in the ventromedial hypothalamus govern mating in both sexes and aggression in males. Cell 153, 896–909.

Yang, C.F., and Shah, N.M. (2014). Representing sex in the brain, one module at a time. Neuron 82, 261–278.

Yang, W., Salil, S.B., and David, M. (2019). *Drosophila* female-specific brain neuron elicits persistent position- and direction-selective male-like social behaviors. bioRxiv, 594960.

Zheng, Z., Lauritzen, J.S., Perlman, E., Robinson, C.G., Nichols, M., Milkie, D., Torrens, O., Price, J., Fisher, C.B., Sharifi, N., et al. (2018). A Complete Electron Microscopy Volume of the Brain of Adult Drosophila melanogaster. Cell 174, 730–743.e722.

Zhou, C., Pan, Y., Robinett, C.C., Meissner, G.W., and Baker, B.S. (2014). Central brain neurons expressing doublesex regulate female receptivity in Drosophila. Neuron 83, 149–163.

Zuiderveld, K. (1994). Contrast limited adaptive histogram equalization. Graphics gems, 474–485.

