## Supplementary figures for "Neurons underlying aggressive actions that are shared by both males and females in *Drosophila*"

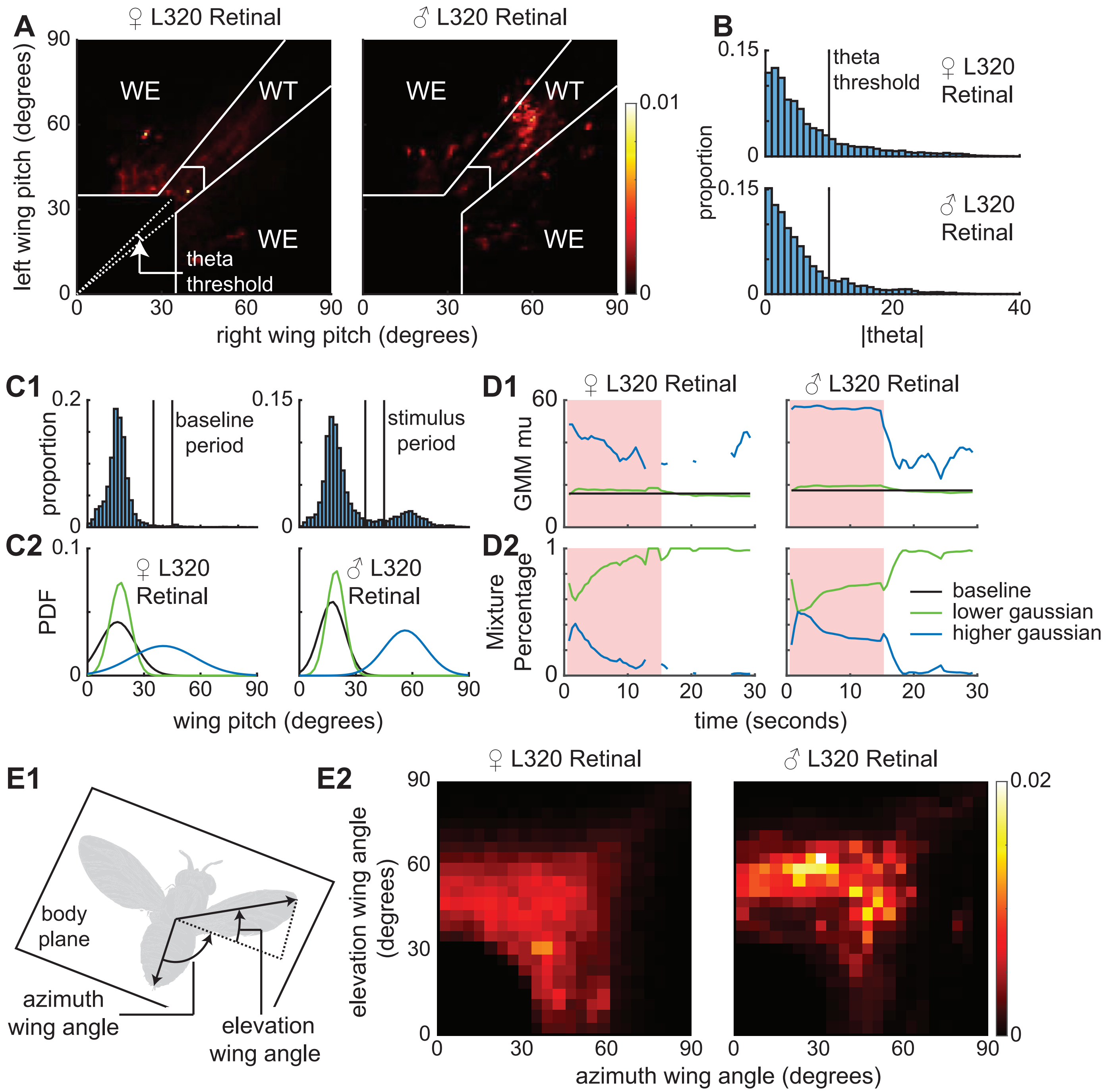


**Figure 1 S1. Method for determining wing extension vs wing threat. A.** The wing pitch space broken down into different wing behavior domains (WT = wing threat and WE = wing extension) based on the wing pitch and angle from the line of unity thresholds. **B.** Probability distribution of the absolute angle the right and left wing pitch makes from the identity line (left pitch = right pitch). The black line represents the threshold used in Figure 1 analysis and shown in **A**. **C1.** Probability distributions of wing pitch for male and female L320 > UASChrimson flies before the stimulus period (**Left**) and during the stimulus period (**Right**) follow a roughly Gaussian and two component Gaussian mixture distribution respectively (**C2**). The two black lines represent the threshold used in the Figure 1 analysis and in panel **A**. **D.** Two component Gaussian mixture model (GMM) fits to 1.5 second (1 s overlap) sliding window wing pitch for female (**Left**) and male (**Right**) flies (see methods). **D1.** The means of the GMM fits with baseline being Gaussian fit to the before light on period shows that flies display both a higher wing pitch and a lower wing pitch throughout the stimulus period. The average lower wing pitch is higher than baseline during the stimulus period and returns to the baseline level after the stimulus is turned off. **D2.** The mixing proportions of the two Gaussians in the GMM show a clear drop in the proportion of higher wing pitch over time for female flies and a slower reduction in this proportion for male flies. **E.** The majority of wing behavior is a combination of an increase in both azimuth and elevation wing angle. **E1.** Schematic illustrating elevation and azimuth wing angle. The body plane is defined as the plane that is orthogonal to the head-thorax-abdomen plane and contains the body axis (see methods). **E2.** Distribution of elevation and azimuth wing angles for female and male L320 > UASChrimson flies during wing threat and wing extension.

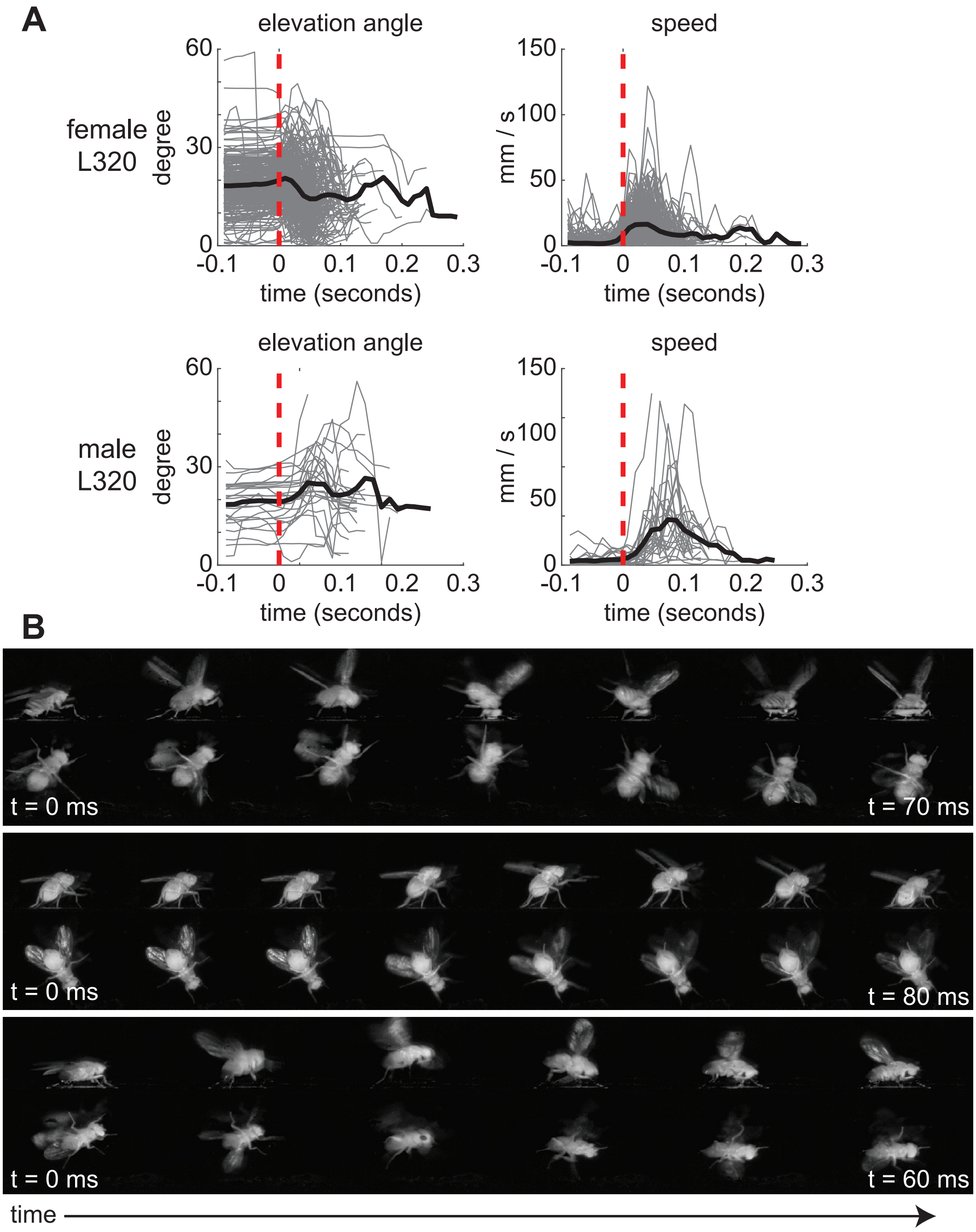


**Figure 1 S2. Characterization of elevation angle and speed underlying thrusting. A.** Elevation angle and speed aligned by manually annotated bouts of high fence used as ground truth (n = 75 female trials, n = 60 male trials). Red dotted line indicates start of a thrust bout. Thrust is characterized by an increase in elevation angle as the fore-legs are raised. At the end of the thrust, flies will snap the forelegs down and drive the body towards the ground, resulting in a drop in the elevation angle. **B.** Three examples of different types of thrusting. **Top:** high intensity female thrust. **Middle:** low intensity female thrust. **Bottom:** high intensity male thrust.

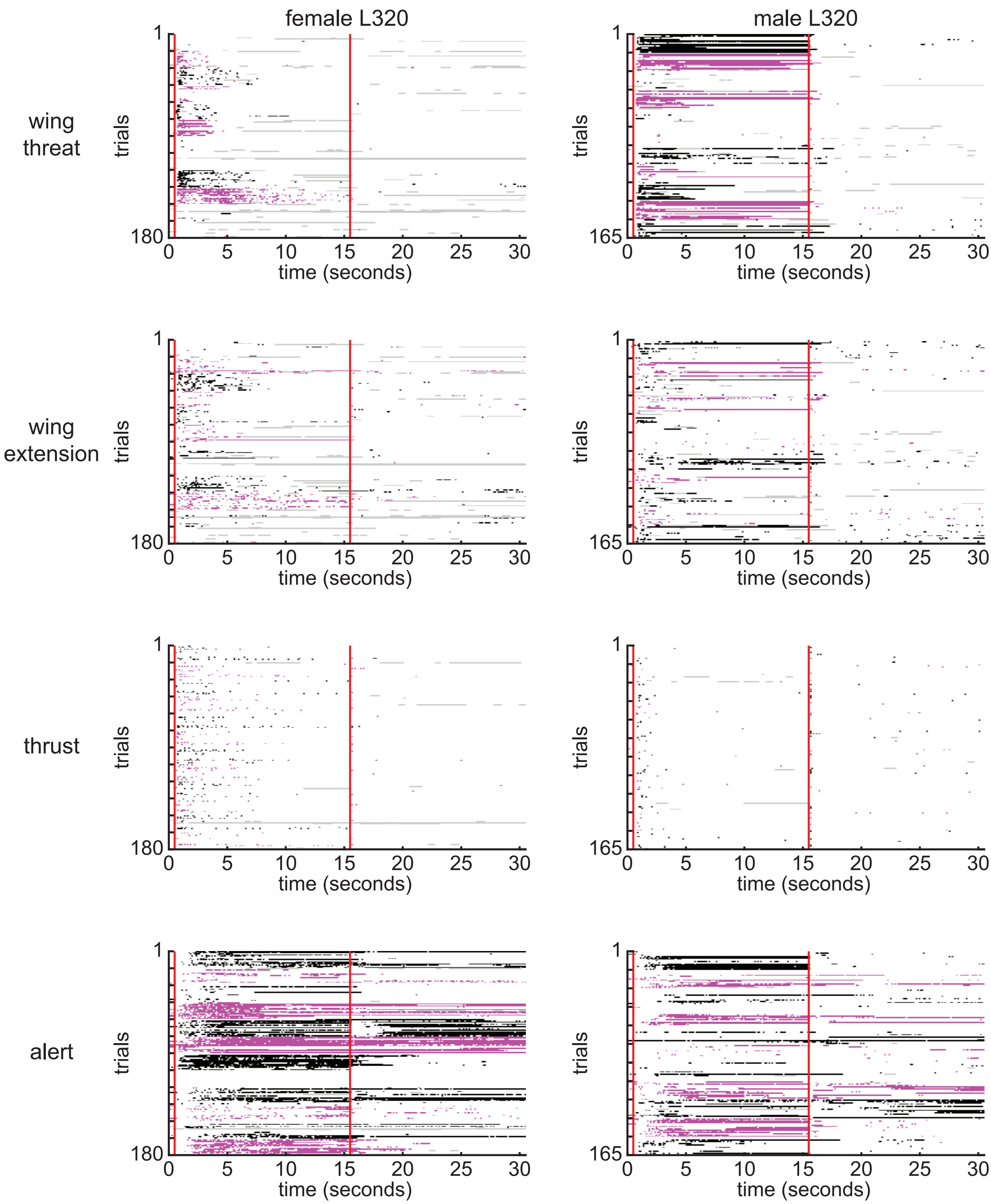


**Figure 1 S3. Single trial ethogram of actions for retinal fed L320 > UASChrimson flies.** Trials grouped by flies (fly 1 = trials 1-15, fly 2 = trials 16-30, etc). Red bars indicate start and end of optogenetic stimulation period. Alternating black and magenta blocks of trials represent different flies. Gray areas indicate >500 ms bouts where there is low confidence in any of the tracked body parts that is used to identify the action.

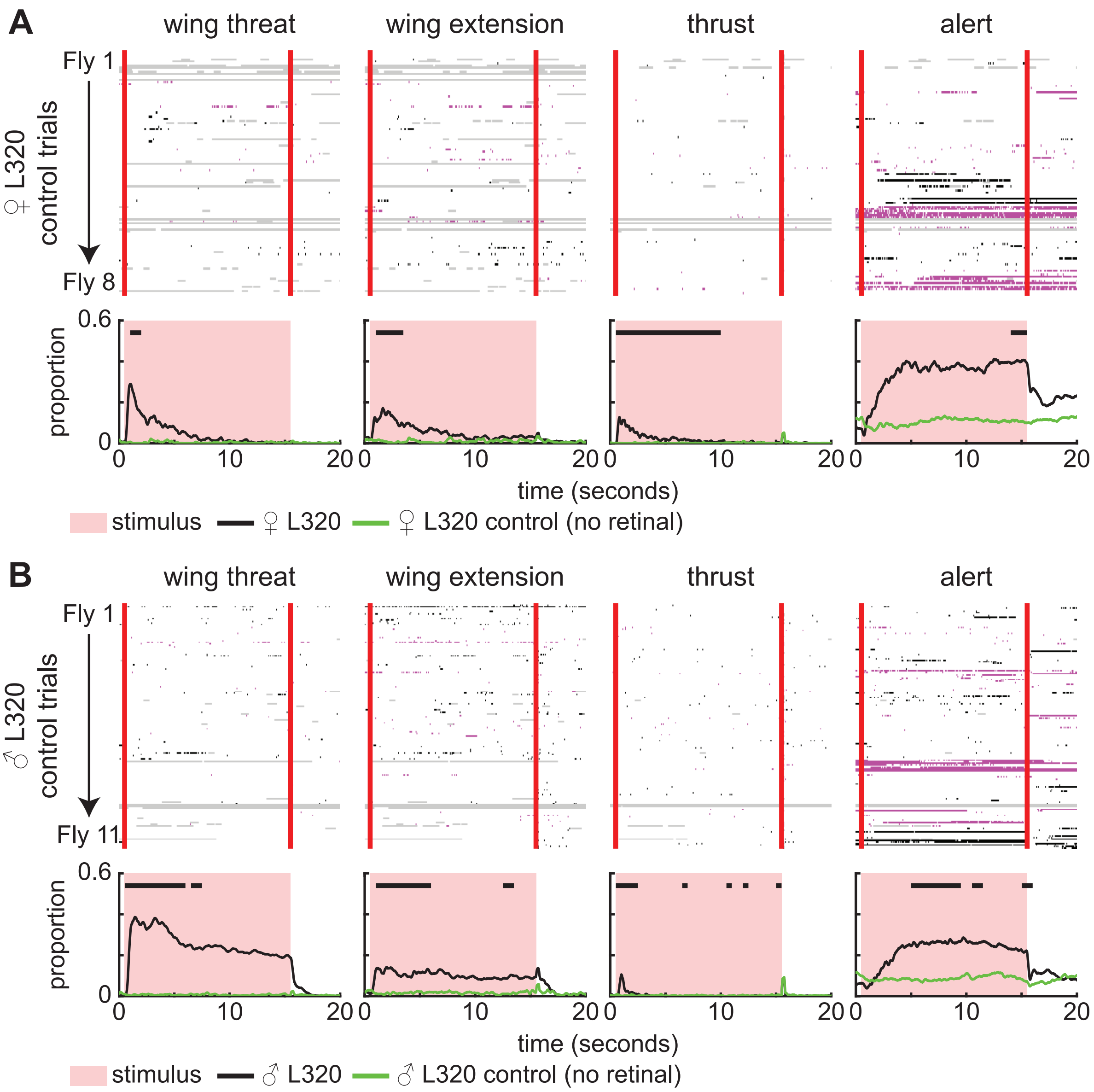


**Figure 1 S4. L320 > UASChrimson flies not fed with retinal do not display aggressive actions. A. Top row:** Single trial ethogram of actions for ♀ L320 > UASChrimson flies not fed with retinal. Trials grouped by flies (fly 1 = trials 1-15, fly 2 = trials 16-30, etc). Red bars indicate start and end of optogenetic stimulation period. Alternating black and magenta blocks of trials represent different flies. Gray areas indicate >500 ms bouts where there is low confidence in any of the tracked body parts that is used to identify the action. **Bottom row:** Proportion of trials where flies are performing each action. An 8.9 mW/cm² 617 nm light is turned on during the stimulus period. (Retinal fed: n=12 flies, Control: n=8 flies; 15 trials/fly). Black bars show time points where there is a significant difference between retinal fed and control flies (Wilcoxon rank sum test p<0.01, see methods). **B.** Same as **A**, but for ♂ L320 > UASChrimson flies. (Retinal fed: n=11 flies; Control: n=11 flies; 15 trials/fly).

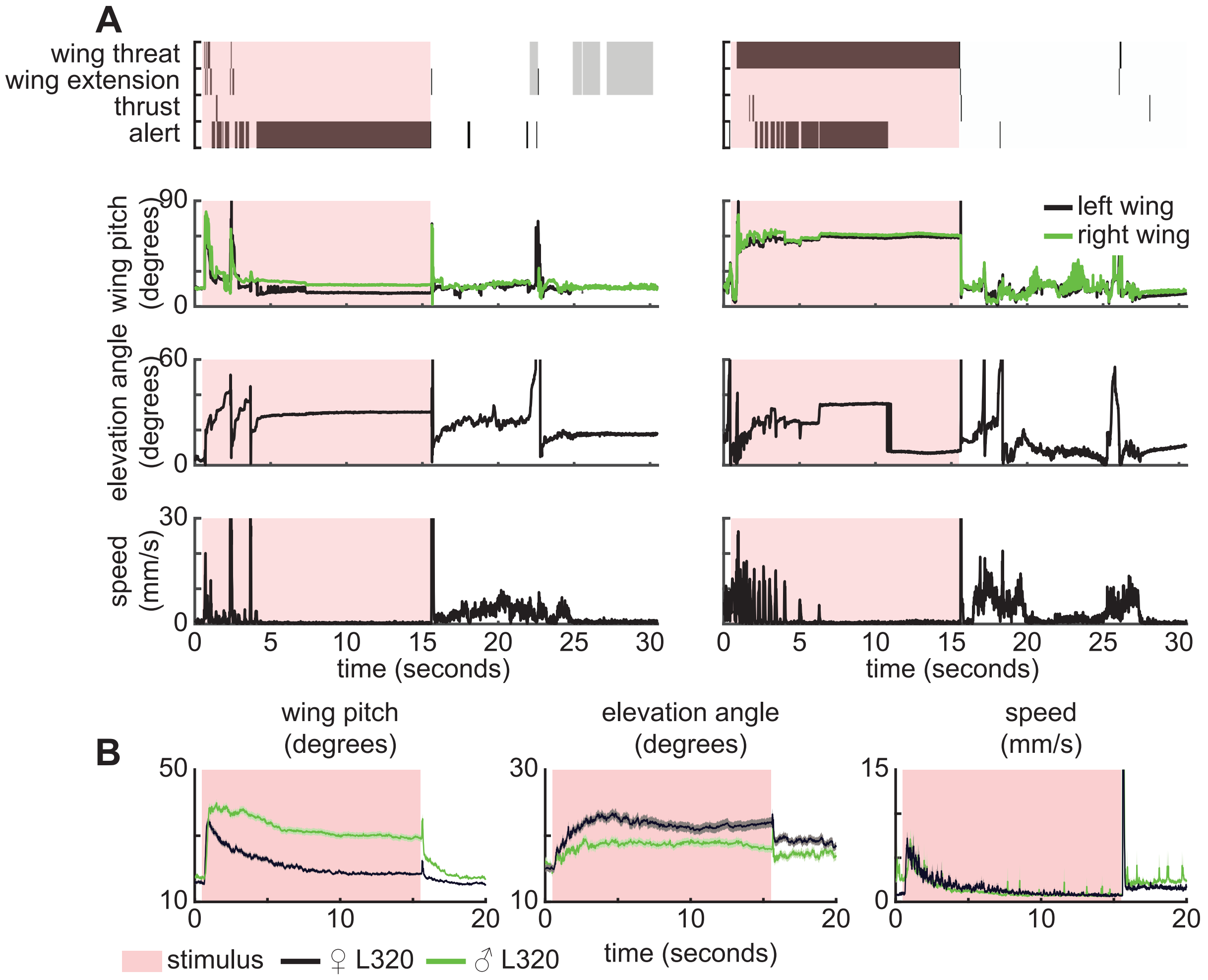


**Figure 1 S5. L320 behavior is stochastic and arises from dynamically changing observables. A.** Sample action ethogram and observables for a single female (left) and male (right) L320 > UASChrimson trial. Gray areas indicate >500 ms bouts where there is low confidence in any of the tracked body parts that are used to identify the action. **B.** Male and female trial averaged wing pitch, speed, and elevation angle over time. Panels show mean +/- standard error of mean.

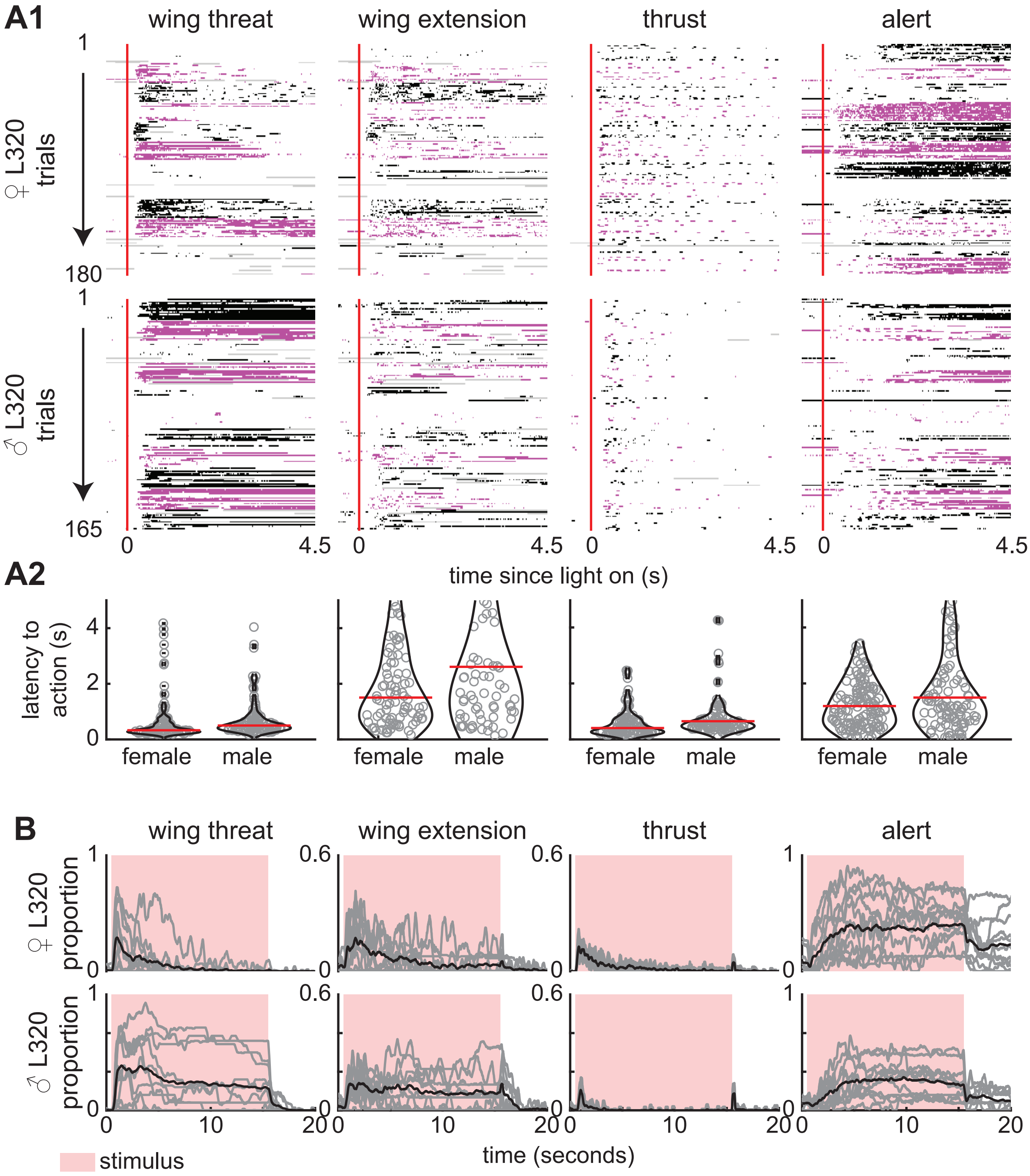


**Figure 1 S6. Optogenetic activation of L320 neurons drives action production at low latency. A1.** Single trial ethogram of actions for male and female L320 > UASChrimson flies for the first 5 seconds. Trials grouped by flies (fly 1 = trials 1-15, fly 2 = trials 16-30, etc). Red line indicates start of optogenetic stimulation period. Alternating black and purple blocks of trials represent different flies. Gray areas indicate frames where any of the tracked body parts used to identify the action had low tracking confidence (see methods). **A2.** Latency from stimulus onset to first instance of each action across all trials. Violin plots show distribution of latency. Red line shows median latency. **B.** Proportion of trials that each male and female L320 > UASChrimson is performing each action as a function of time. Each grey line is a fly and the black line is the average across all flies. An 8.9 mW/cm² 617 nm light is turned on during the stimulus period. (Retinal fed: n=12 female flies, n=11 male flies; 15 trials/fly).

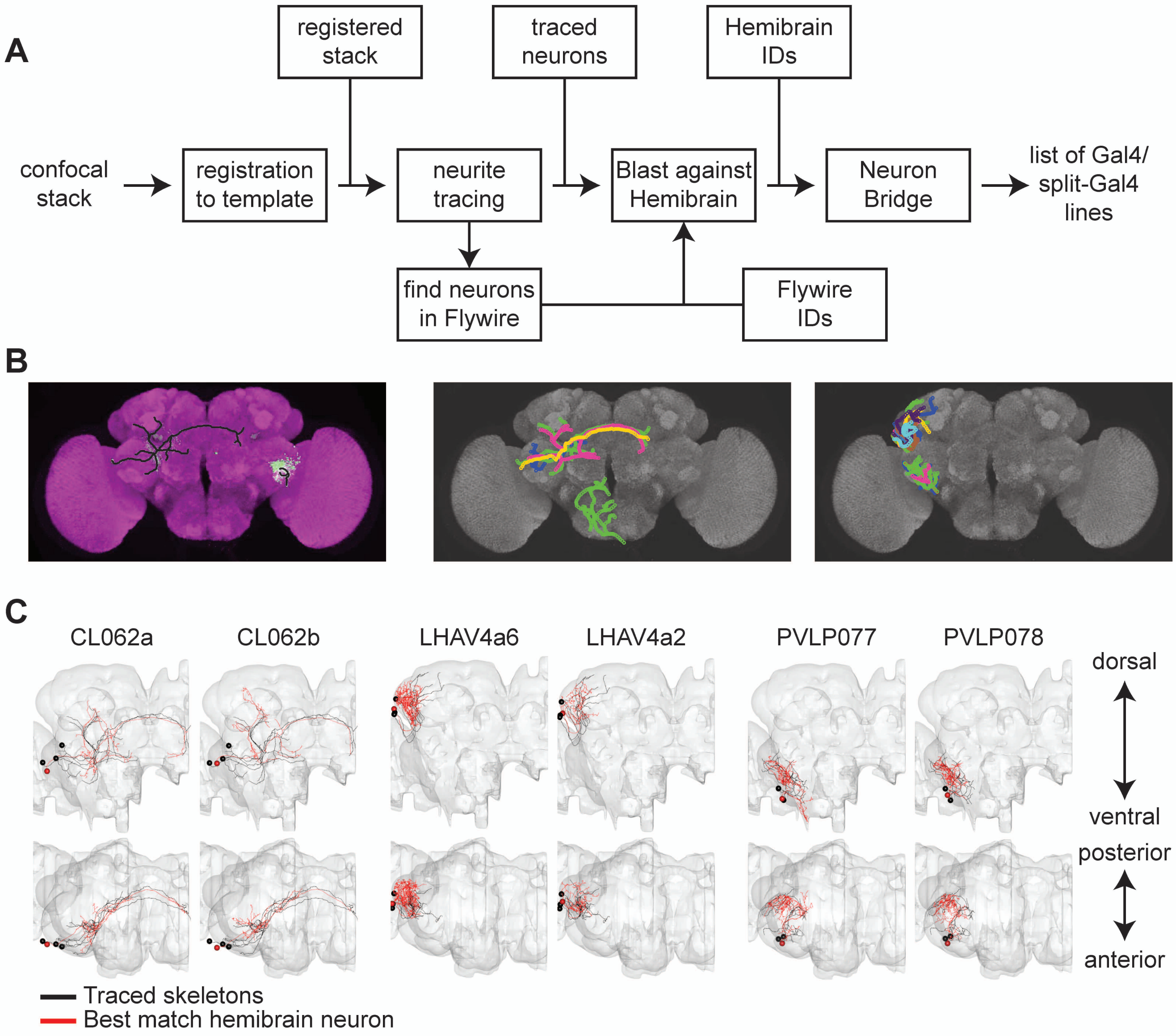


**Figure 2 S1. Finding hemibrain and flywire IDs for L320 neurons. A.** Analysis pipeline to find hemibrain and flywire IDs for neurons labeled by L320. **B. Left:** Sample MCFO brain registered to JFRC2 with neurons in green and JFRC2 template brain in purple. The traced skeletons for neurons are shown in black. **Middle:** All skeletonized right hemisphere neurons in the clamp and the single descending neuron. **Right:** Same as middle panel, but for neurons in the anterior lateral horn and posterior ventro-lateral protocerebrum. Each color represents a different neuron trace. **C.** Top matched hemibrain classes for each cluster of neurons labeled by L320. Sample traced skeletons from multicolor flipout are shown in black. The top matched hemibrain neurons based on nblast and visual inspection are shown in red.

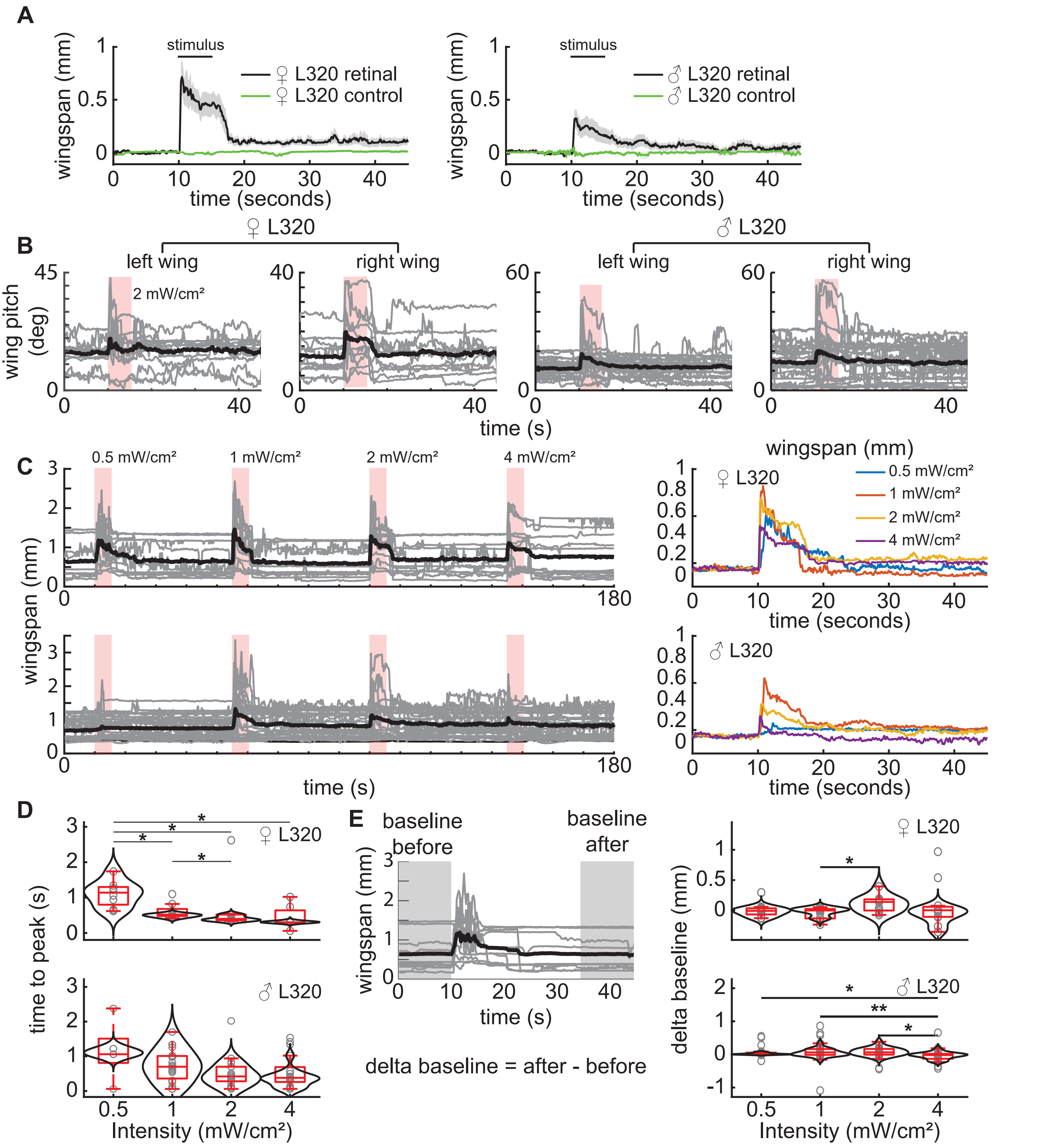


**Figure 3 S1. Optogenetic stimulation in head fixed open cuticle flies is capable of driving wing behavior. A.** L320 > UASChrimson flies not fed with retinal do not display behavior. (control Females: n=20 trials across 5 flies, control Males: n=20 trials across 5 flies). Panels show mean +/- standard error of mean. **B.** Single trial left and right wing pitch of L320 > UASChrimson flies in a dissected head fixed track-ball setup. A 2 mW/cm² 617 nm light is delivered during the 5 second stimulus period. (Females: n=11 trials across 5 flies, Males: n=26 trials across 7 flies) **C.** Same as **B**, but for wingspan. Each trial comprises of 4 increasing light intensities with a 30 second rest period in between. **Left**: All trials across female (**Top**) and male (**Bottom**) flies. **Right**: Mean wingspan averaged across trials based on stimulus intensity for females (**Top**) and males (**Bottom**). Trials were baseline subtracted and aligned by 10 seconds before stimulus onset. **D.** Latency to the first the peak in wingspan after stimulus light on for female (**Top**) and male (**Bottom**) flies. Trials where the wingspan did not change were not considered. (Wilcoxon rank sum test * $p<0.05$, ** $p<0.01$) **E.** Left: Schematic illustrating the change in baseline wingspan in the time period 10 seconds before stimulus onset and 20 to 30 seconds after stimulus offset. Right: Change in baseline wingspan for female (**Top**) and male (**Bottom**) flies. (Wilcoxon signed-rank test * $p<0.05$, ** $p<0.01$)

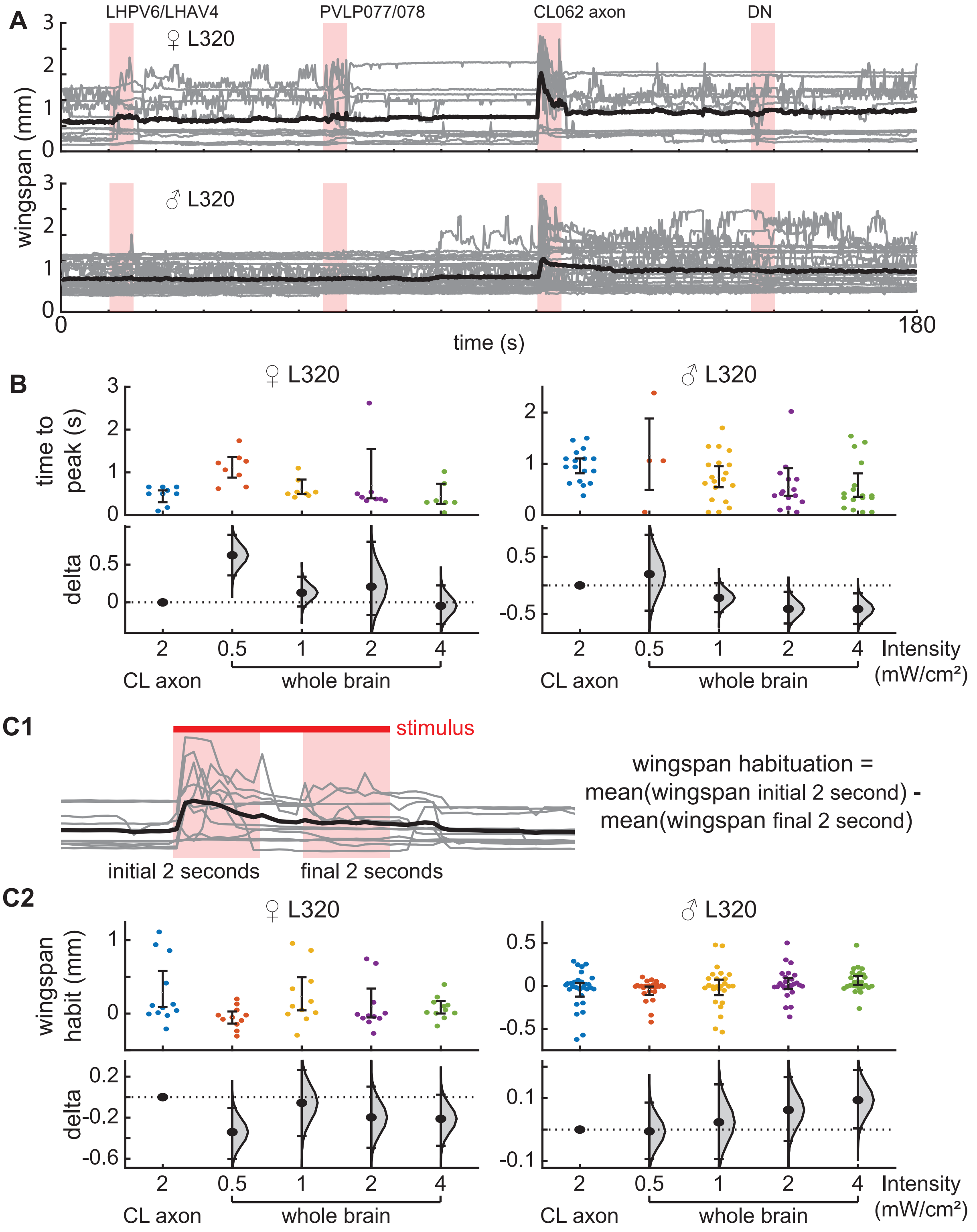


**Figure 3 S2. Activation of only CL062 axons drives wing behavior. The level of this behavior is similar to the activation of all L320-split neurons at half the light intensity. A.** Wingspan of UASChrimson>L320 split-gal4 flies in a dissected head fixed trackball setup. Each trial comprises of sequentially activating the LH neurons, the PVLP neurons, the CL062 axons, and finally the DN. Neurons are activated using a 2 mW/cm² 617 nm light through a DMD projector. There is a 30 second rest period between each stimulus application. All trials across female (**Top**) and male (**Bottom**) flies (Males: n=29 trials across 7 flies, Females: n=12 trials across 5 flies). **B.** Latency to the peak of wingspan after stimulus light on for female (**Left**) and male (**Right**) when the CL062 axons were targeted with a 2 mW/cm² 617 nm as compared to optogenetic stimulation of all neurons at different intensities. Trials where the wingspan did not peak. **C1.** Definition of wingspan habituation as the difference in mean wingspan during the first 2 seconds of each stimulus bout minus the mean wingspan during the last 2 seconds of each stimulus bout. **C2.** Wingspan habituation for female (Left) and male (Right) when the CL062 axons were targeted with a 2 mW/cm² 617 as compared to optogenetic stimulation of all neurons at different intensities.

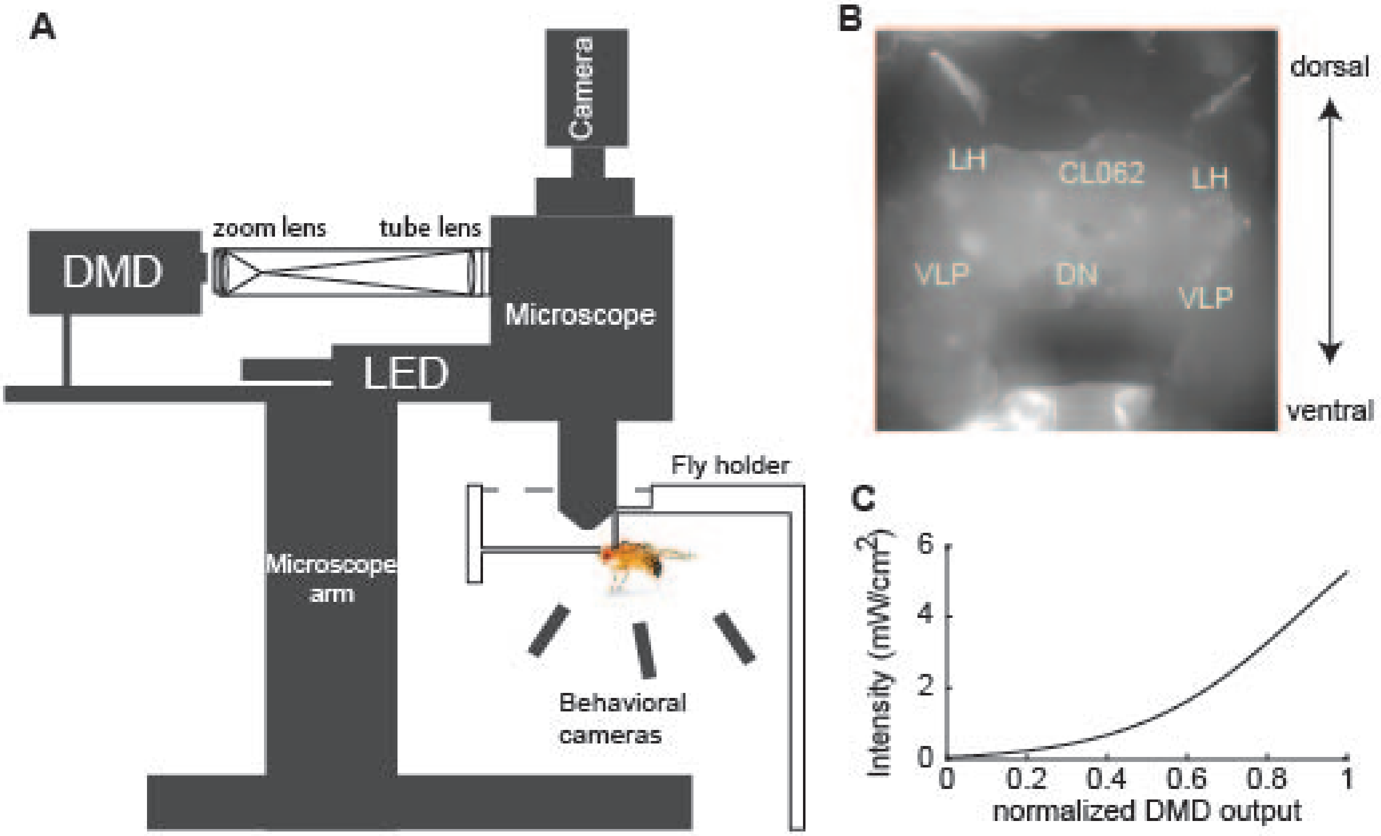


**Figure 3 S3. Digital micromirror device (DMD) experimental setup. A.** Hardware setup. A achromatic doublet zoom lens (AC254-035-A-ML) and tube lens was used to bring the projector image to the microscope. **B.** Max projection image of neurons under the microscope. C. Intensity calibration curve for 617 nm.

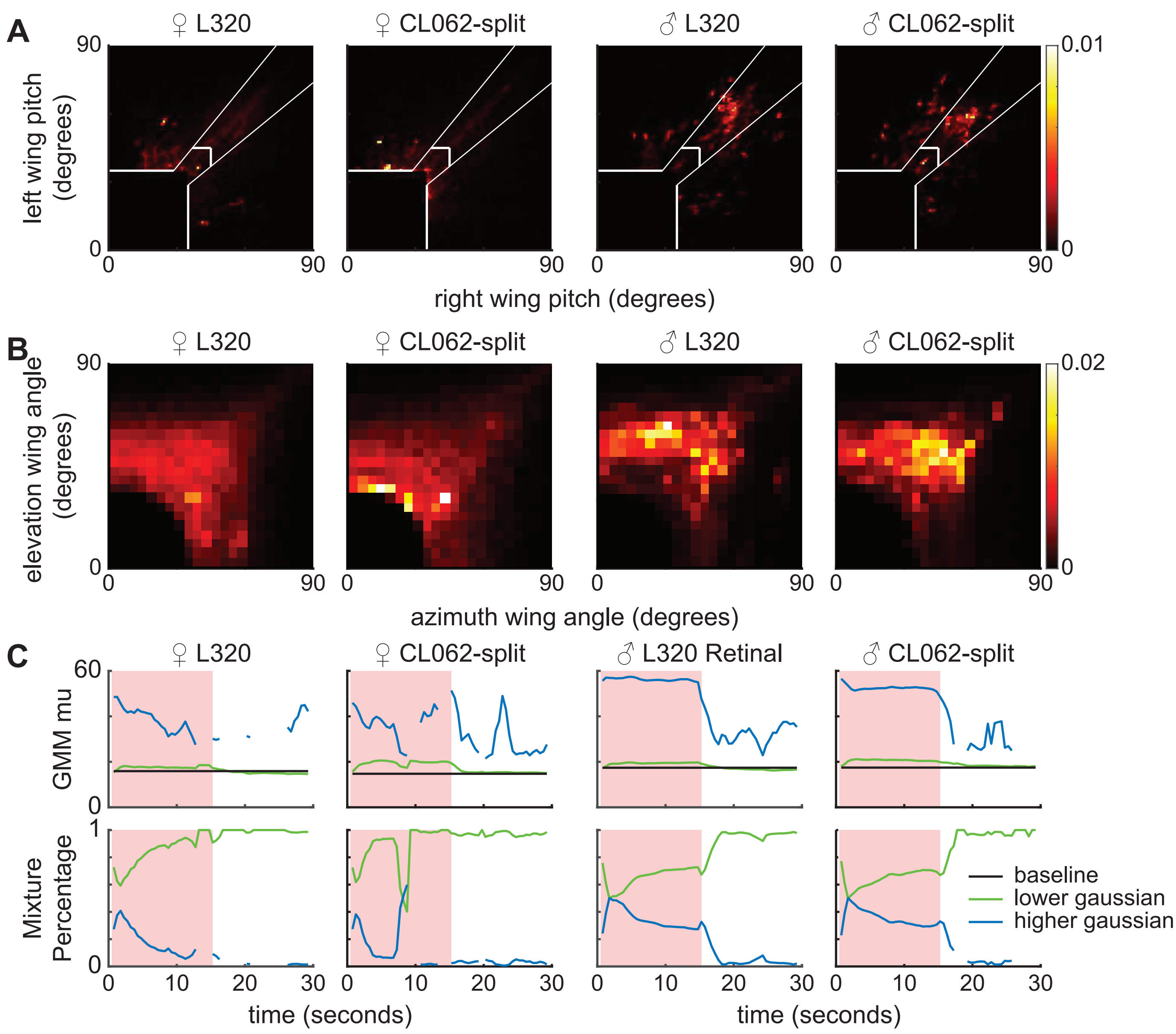


**Figure 4 S1. CL062-split flies exhibit similar wing activity as L320 flies. A.** The wing pitch space broken down into different wing behavior domains based on the wing pitch and angle from the line of unity thresholds shows similar distribution of wing position for male L320 > UASChrimson and CL062-split > UASChrimson flies. Female flies of each genotype also show similar distribution. **B.** Distribution of elevation and azimuth wing angles during wing threat and wing extension again show sex matched similarity in distribution across genotypes. **C.** Two component Gaussian mixture model (GMM) fits to 1.5 second (1 s overlap) sliding window wing pitch for female and male L320 > UASChrimson and CL062-split > UASChrimson flies show similar GMM means and temporal dynamics or mixing proportions.

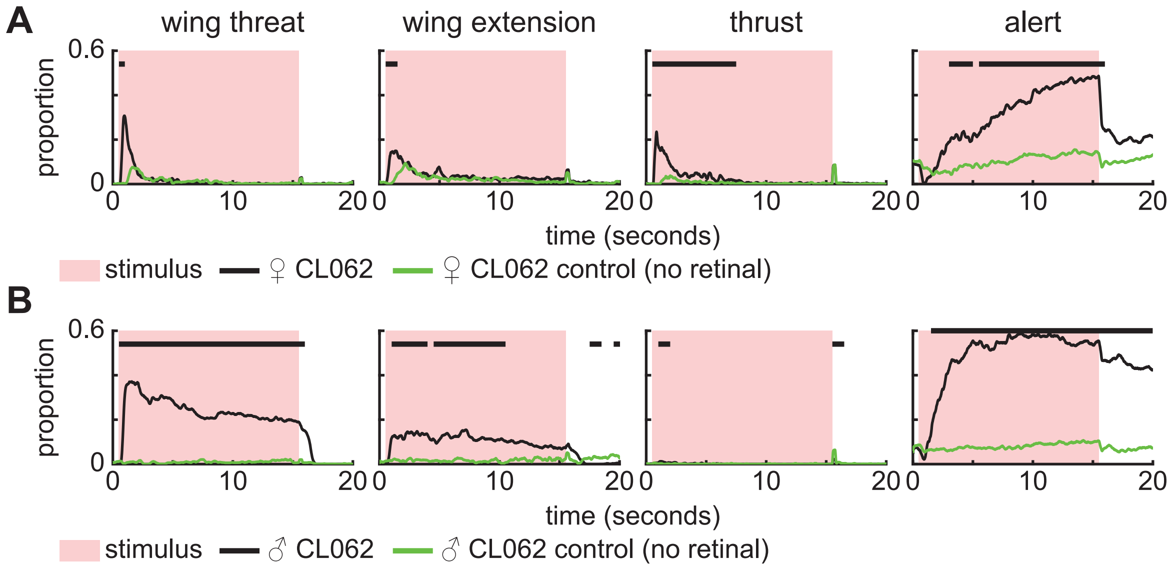


**Figure 4 S2. CL062-split > UASChrimson flies not fed with retinal do not display aggressive actions. A.** Proportion of trials where ♀ CL062-split > UASChrimson flies are performing each action. An 8.9 mW/cm² 617 nm light is turned on during the stimulus period. (Retinal fed: n=11 flies, Control: n=12 flies; 15 trials/fly). Black bars show time points where there is a significant difference between retinal fed and control flies (Wilcoxon rank sum test p<0.01, see methods). **B.** Same as **A**, but for ♂ CL062-split > UASChrimson flies. (Retinal fed: n=11 flies; Control: n=11 flies; 15 trials/fly).

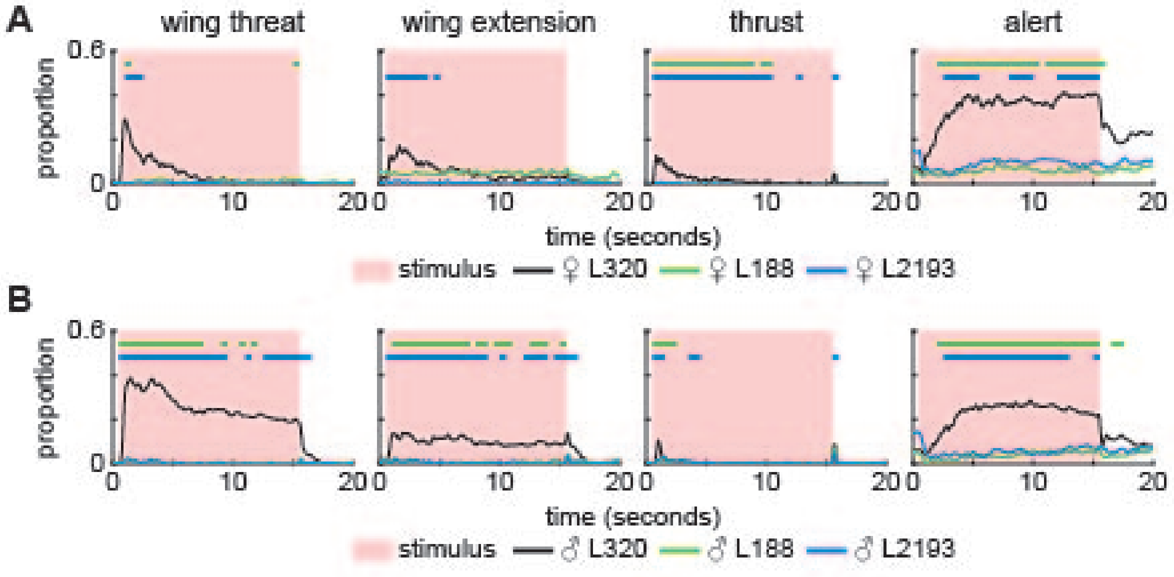


**Figure 4 S3. LHAV4a and LHPV6a1/3 does not drive aggressive actions. A.** L188 split-GAL4 and L2193 split-GAL4 labels LHPV6a1 and LHPV6a3 neurons respectively. Shown are the proportion of trials where female L188 split-GAL4 > UASChrimson and L2193 split-GAL4 > UASChrimson flies are performing each action. An 8.9 mW/cm² 617 nm light is turned on during the stimulus period. (Retinal fed: n=11 L188 flies, n=10 L2193 flies; 15 trials/fly). Green and blue bars show time points where there is a significant difference between L320 > UASChrimson and L188 > UASChrimson and L2193 > UASChrimson flies respectively (Wilcoxon rank sum test p<0.01, see methods).**B.** Same as **A**, but for male flies. (Retinal fed: n=9 L188 flies, n=10 L2193 flies; 15 trials/fly)

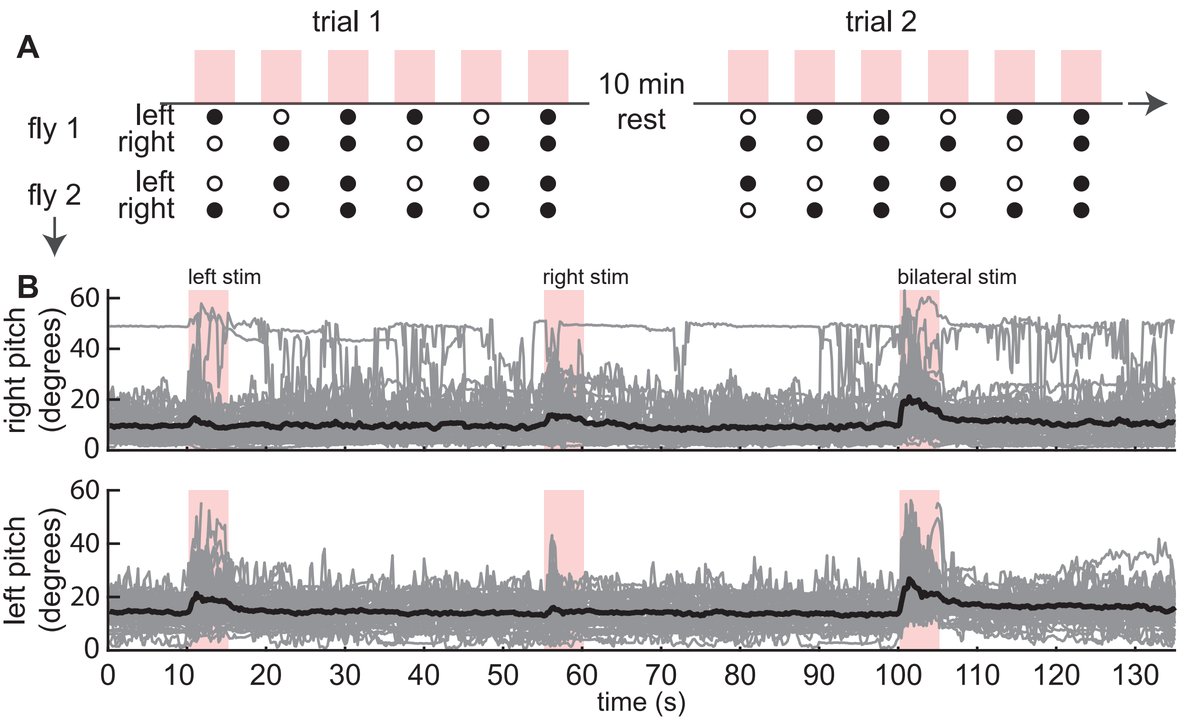


**Figure 5 S1. Single trial motor elements during unilateral and bilateral stimulation. A.** Experimental setup. Each trial block involved two rounds of unilateral stimulation followed by bilateral stimulation. We alternated whether the left or right hemisphere was stimulated first across trials and for different flies. **B.** Wing pitches after left, right, and bilateral stimulation. For readability, we ordered the stimulus in each trial blocks such that the left stimulus occurs first. (Females: n=23 trials across 5 L320 > UAS-Chrimson flies).

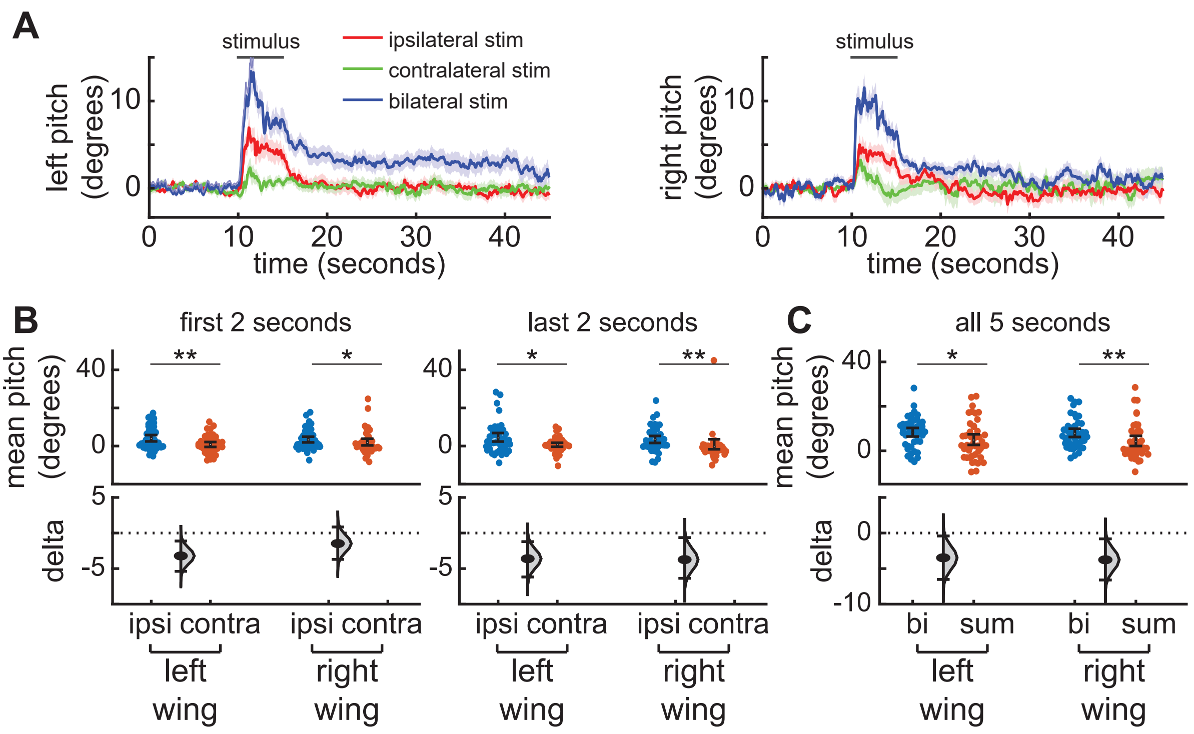


**Figure 5 S2. Unilateral activation of CL062 neurons shows that ipsilateral and contralateral wings threats are controlled differentially. A.** Baseline subtracted left and right wing pitch after ipsilateral, contralateral, and bilateral stimulation shows a fast increase in bilateral wing pitch followed by higher ipsilateral wing pitch. (Females: n=23 trials across 5 L320 > UASChrimson flies). Panels show mean +/- standard error of mean. Neurons are activated using a 2 mW/cm² 617 nm light delivered at the neuron focal plane. **B. Left:** The baseline subtracted mean left and right wing pitch during the first 2 seconds of ipsilateral stimulus is higher than that of contralateral stimulus despite bilateral wing pitch changes. **Right:** The baseline subtracted mean left and right wing pitch during the first 2 seconds of ipsilateral stimulus is higher than that of the contralateral stimulus. Only the right wing pitch is significantly higher. **C.** The baseline subtracted mean pitch during the entire 5 second bilateral stimulation period is significantly higher than the sum of the mean left and right pitch during the stimulus period (Wilcoxon rank sum test * $p<0.05$, ** $p<0.01$).

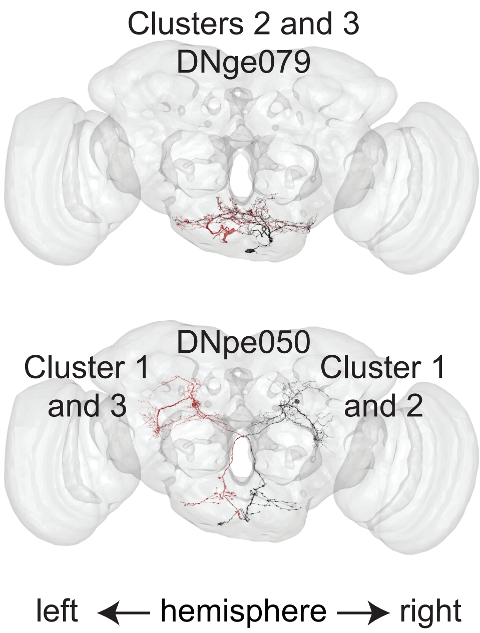


**Figure 6 S1. Trends in DN organization based on connections to CL062 neurons.** **Top:** Two pairs of DNs in the Gnathal Ganglia are strongly connected to cluster 2 and 3 CL062 neurons. **Bottom left:** DNpe050 is strongly connected to ipsilateral clusters (2 and 3) of CL062 neuron and to the bilateral cluster of Cl062 neuron (1). **Bottom right:** The pair of fruitless descending neurons predicted to be strongly connected to cluster 1 Cl062 neurons.

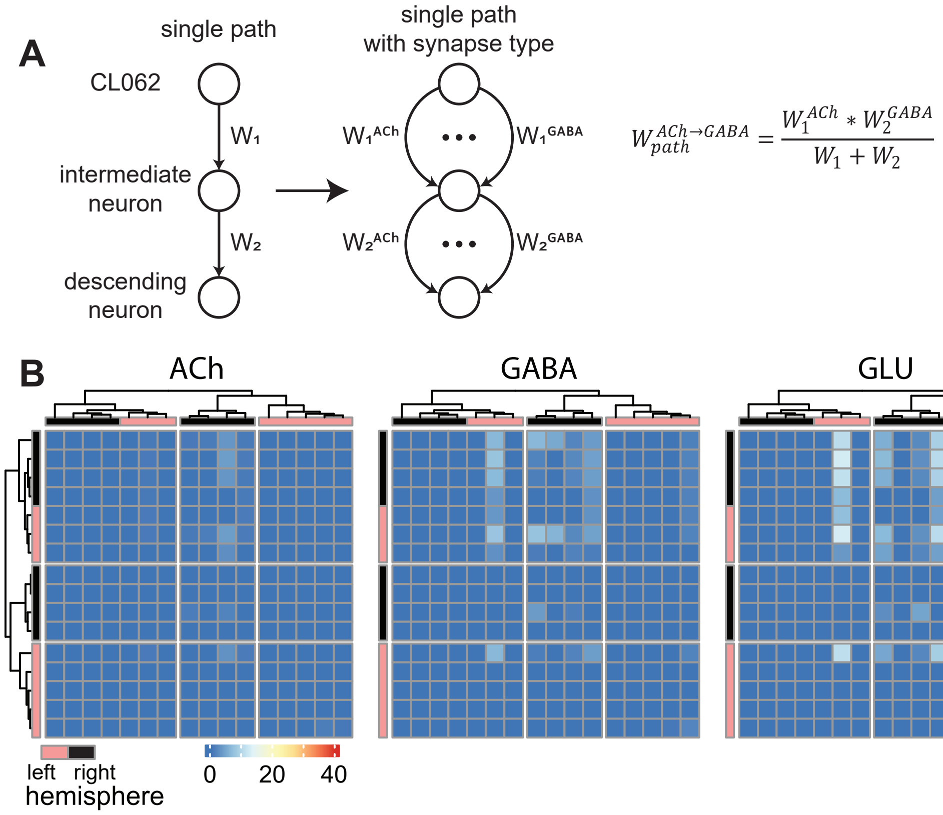


**Figure 6 S2. CL062 makes limited first and second order recurrent connections. A.** Schematic of decomposing paths based on neurotransmitter type. **B.** Neurotransmitter pathways from CL062 neurons to each other show weak and sparse GABA and Glutamate pathways between CL062 neurons. CL062 clusters based on Figure 6.

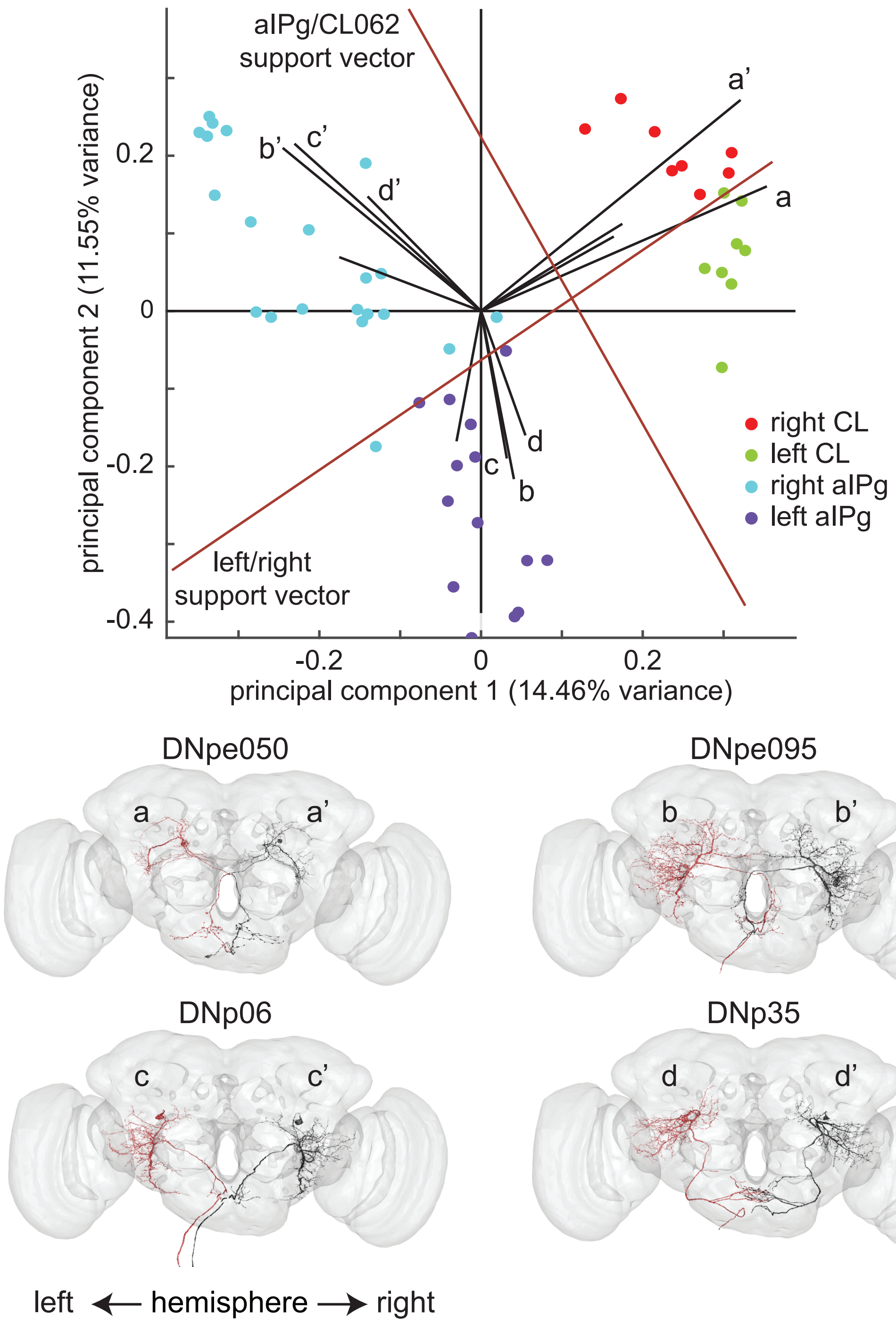


**Figure 7 S1. Identification of potentially important DNs that contribute to the parallel descending pathways of CL062 and aIPg neurons. Top.** Biplot of the first two principal components (PCs) of L2 normalized connections to descending neurons from all CL, aIPg, and pC1d/e neurons. Top four loading vectors in the direction of CL062, left aIPg, and right aIPg neuron clusters show descending neurons (DNs) with high contributions to the separation of these clusters in the first two PC space. **Bottom.** Of these 12 DN loadings, there are 4 pairs of DNs. Based on the direction of their loadings, these neurons are most strongly connected with ipsilateral CL and aIPg neurons respectively.

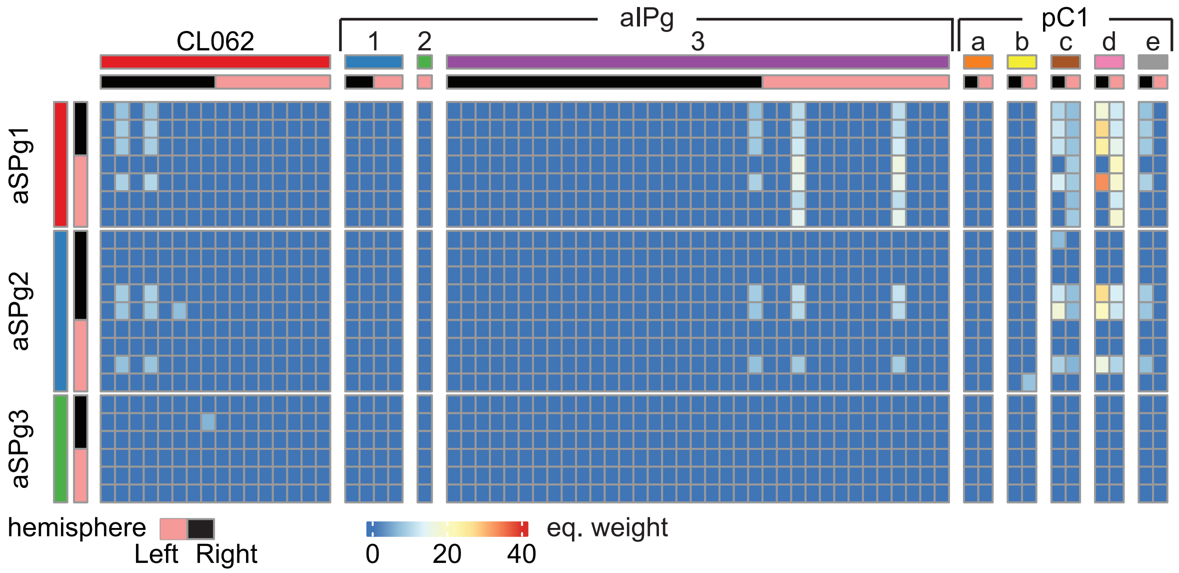


**Figure 8 S1. aSPg makes connections to pC1 and sparse connections to CL062.** Matrix (row -> column) showing equivalent weight from aSPg to CL062, aIPg, and pC1 neurons. Pathways include both direct connections and one layer removed indirect connections via an intermediate neuron.
